# Variation in reflectance spectroscopy of European beech leaves captures phenology and biological hierarchies despite measurement uncertainties

**DOI:** 10.1101/2021.03.09.434578

**Authors:** Fanny Petibon, Ewa A. Czyż, Giulia Ghielmetti, Andreas Hueni, Mathias Kneubühler, Michael E. Schaepman, Meredith C. Schuman

## Abstract

The measurement of leaf optical properties (LOP) using reflectance and scattering properties of light allows a continuous, time-resolved, and rapid characterization of many species traits including water status, chemical composition, and leaf structure. Variation in trait values expressed by individuals result from a combination of biological and environmental variations. Such species trait variations are increasingly recognized as drivers and responses of biodiversity and ecosystem properties. However, little has been done to comprehensively characterize or monitor such variation using leaf reflectance, where emphasis is more often on species average values. Furthermore, although a variety of platforms and protocols exist for the estimation of leaf reflectance, there is neither a standard method, nor a best practise of treating measurement uncertainty which has yet been collectively adopted. In this study, we investigate what level of uncertainty can be accepted when measuring leaf reflectance while ensuring the detection of species trait variation at several levels: within individuals, over time, between individuals, and between populations. As a study species, we use an economically and ecologically important dominant European tree species, namely *Fagus sylvatica*. We first use fabrics as standard material to quantify the measurement uncertainties associated with leaf clip (0.0001 to 0.4 reflectance units) and integrating sphere measurements (0.0001 to 0.01 reflectance units) via error propagation. We then quantify spectrally resolved variation in reflectance from *F. sylvatica* leaves. We show that the measurement uncertainty associated with leaf reflectance, estimated using a field spectroradiometer with attached leaf clip, represents on average a small portion of the spectral variation within a single individual sampled over time (2.7 ± 1.7%), or between individuals (1.5 ± 1.3% or 3.4 ± 1.7%, respectively) in a set of monitored *F. sylvatica* trees located in Swiss and French forests. In all forests, the spectral variation between individuals exceeded the spectral variation of a single individual measured within one week. However, measurements of variation within an individual at different canopy positions over time indicate that sampling design (e.g., standardized sampling, and sample size) strongly impacts our ability to measure between-individual variation. We suggest best practice approaches towards a standardized protocol to allow for rigorous quantification of species trait variation using leaf reflectance.

**Highlights:**

- We partition biological variation from measurement uncertainty for leaf spectra.
- Measurement uncertainty represents ca. 3% of spectral variation among beech trees.
- Biological variation within an individual increases by 80% as leaves mature.
- Maxima of uncertainty correspond to maxima of biological variation (water content).
- We recommend procedures to quantify biological variation in spectral measurements.

## 1. Introduction

Monitoring biodiversity across biological scales – from genetic diversity to functional trait diversity – is critical to assess biodiversity loss, shifts, and resilience under increasingly rapid global change (Hansen et al., 2001; Oehri et al., 2017). The Group on Earth Observations Biodiversity Observation Network (GEO BON) developed Essential Biodiversity Variables (EBVs) for standardized and harmonized assessment and reporting of biodiversity change (Pereira, 2013). Among the six classes of EBVs, ‘species traits’ encompasses all within-species features (e.g., phenological, physiological) that can be measured at the individual level and that allow for the monitoring of intraspecific variation across space and time (Kissling et al., 2018; Pereira, 2013). Intraspecific variation reflects the breadth of plant functional trait attributes expressed by individuals within a species as a result of genetic diversity and phenotypic plasticity in response to environmental factors (Albert et al., 2011). Greater intraspecific variation promotes species coexistence in a more diverse range of environments (Bolnik et al., 2011; Sides et al., 2014). Monitoring species trait variation offers the promise to better understand the contribution of intraspecific variation to biodiversity, ecosystem properties, and species resilience under global change (Kissling et al., 2018; Violle et al., 2007). However, to date, no species traits EBVs can be measured from space (Kissling et al., 2018; Pettorelli et al., 2016). Specific leaf area is a morphological trait having the greatest potential to remotely sense species traits variation (Ali et al. 2017, Pettorelli et al., 2016).

Leaf optical properties (LOP) describe how leaves reflect, absorb, and transmit light (Jacquemoud and Ustin, 2019a). As chemical and morphological features alter the way leaves interact with light, LOP integrate variation in these properties, which include traits that are indicators of plant ecophysiology and performance. An understanding of LOP allows us to interpret the leaf reflectance measurements retrieved from remote sensing platforms for monitoring the variation of LOP across space and time, allowing for a simultaneous and rapid characterization of several specific functional plant traits (Jacquemoud and Ustin, 2019a).

Numerous studies have demonstrated the potential of remote sensing to monitor functional plant traits at different biological scales (Cavender-Bares et al., 2017, Meireles et al., 2020, Schneider et al., 2017) and have developed biodiversity metrics based on the spectral variation of LOP (Laliberté et al., 2020; Meireles et al. 2020; Schweiger et al., 2018; Williams et al., 2020). However, because of a lack of available data or as a result of concerns that such a level of detail will be neither generalizable nor scalable, very few studies attempted to derive a comprehensive metric for species trait variation or to discern such variation from measurement uncertainty (Cavender-Bares et al., 2016; Čepl et al., 2018; Czyż et al., 2020; Garcia-Verdugo et al., 2010; Madritch et al., 2014; Santiso and Rtuerto, 2015).

Field spectroradiometers are widely used to estimate leaf reflectance at the individual level (Jacquemoud and Ustin, 2019b) and to compare with measurements from airborne optical sensors (Hueni and Bialek, 2017). Coupled with a leaf sampling device and standardized light source, they allow leaf reflectance measurements to be taken independently of environmental conditions and thus with an expected high accuracy and repeatability. The portability of the field spectroradiometer and leaf sampling device allows for non-destructive *in-situ* measurements. While leaf clip (LC) devices measure in a bi-directional view-illumination geometry, integrating sphere (IS) devices integrate the reflected (or transmitted) light over a full hemisphere, which reduces anisotropic directional reflectance (or transmittance) behavior (c.f., Schaepman-Strub et al., 2006 for terminology). For this reason, measurements with an IS are often considered more repeatable and comparable, and thus preferred when sampling conditions allow (Milton, 2009). However, information about anisotropic properties of LOP, which may be important, is lost with IS measurements. Comparison of LC and IS measurements of the same standard or leaf material revealed systematic differences in reflectance retrieved from both types of measurement, although variation among IS measurements was found to be smaller (Hovi et al., 2018; Lukeš, 2017; Potůčková et al., 2016). In this respect, Potůčková et al. (2016), among others, encouraged the standardization of measurement procedures to improve the comparability of spectral measurements across measurement campaigns and within open access spectral librairies.

Nevertheless, the diversity in instrumentation and measurement protocols remains, and the systematic characterization of measurement uncertainty associated with particular protocols is rarely reported.

Measurement uncertainty is inherent to any optical measurement and characterizes the dispersion in reflectance (or absorbtance, or transmittance) that could reasonably be attributed to the target (JCGM, 2008). The optical properties of the target itself, the characteristics of the spectroradiometer and external factors, can impact the dispersion in reflectance. Several studies exist to quantify and minimize measurement uncertainty in spectroradiometric measurements, as well as in (needle) leaf optical properties measurements (Forsström et al., 2021; Helder et al., 2012; Schaepman and Dangel, 2000; Yanez-Rausell et al., 2014a,b). In this study, we investigate to which extent LOP, and particularly the uncertainties associated with leaf reflectance measurements, permit the detection of species traits. We hypothesize (1) that measurement uncertainty depends on the plant sampling device and anisotropic properties of the target, but (2) remains negligible compared to the total variation in spectra measured from different biological samples; and (3) that variation in LOP increases at increasing levels of biological organization from individuals to populations. Lastly, we hypothesize (4) that sampling time, size, and location contribute to the variation in LOP and alter our ability to accurately assess variation between individuals and populations (point 3).

To evaluate our hypotheses, we first consider the spectral reflectance of a set of fabrics with various degrees of anisotropy measured with a field spectroradiometer coupled with either a leaf clip or an integrating sphere. We calculate the measurement uncertainty associated with each material-sampling device pair. We then assess measurement uncertainty and variation at several levels of biological organisation within three datasets comprising the leaf reflectance of *Fagus sylvatica* individuals from several European forests (**Fig. 1**) which is chosen as a dominant tree of economic and ecological importance, having an uncertain future under global change (Brun et al., 2020).

**Figure 1.**
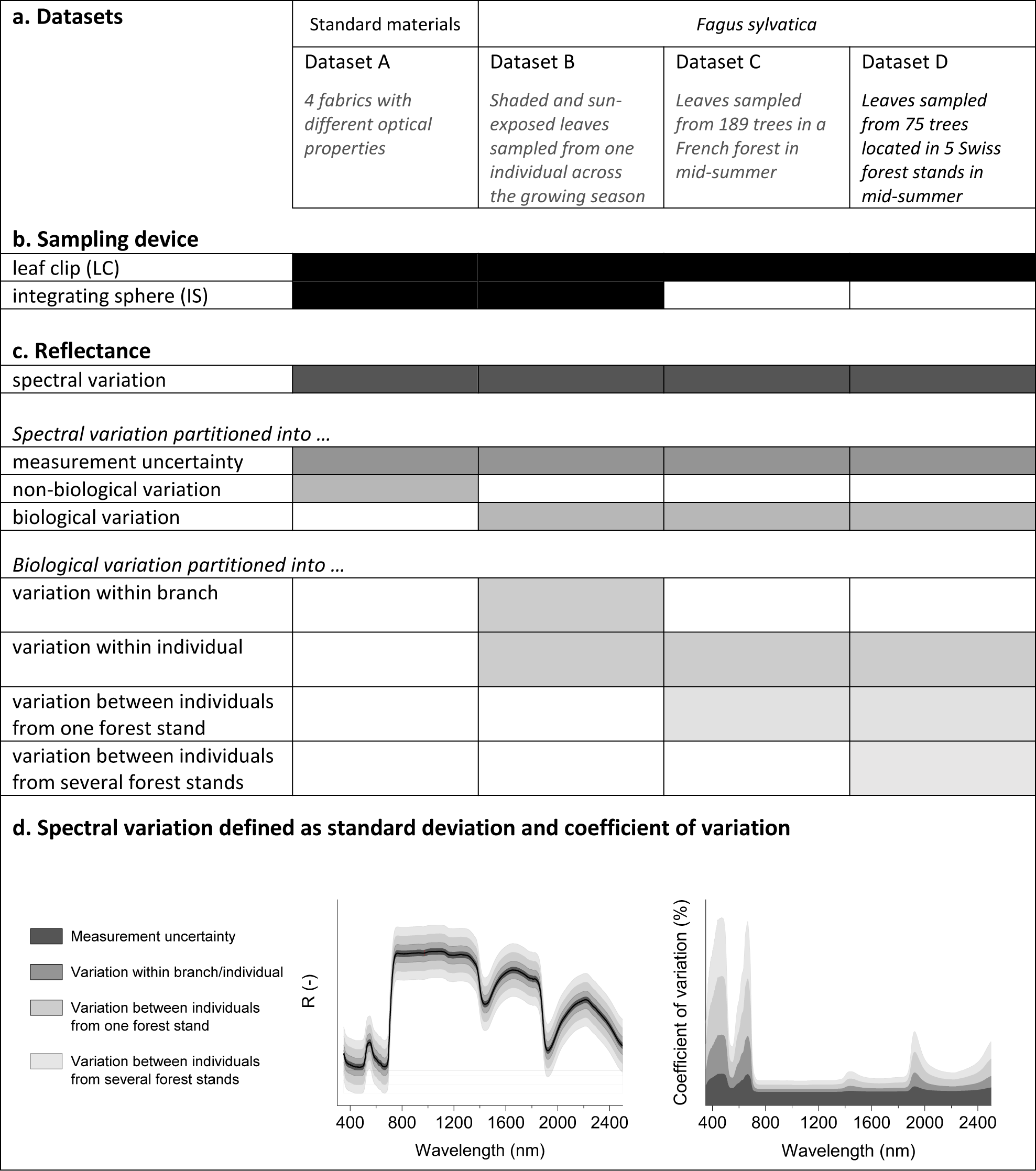
Conceptual overview of the study. The study includes four datasets which allow to assess different sources of variation in optical properties measurements made with a field spectroradiometer and two types of sampling devices, i.e., a leaf clip and an integrating sphere (b). Dataset A is composed of reflectance of a set of fabrics used as standard materials to estimate the measurement uncertainty associated with a sampling device. Sources of uncertainty originate from sensor type, experimental conditions (moisture, temperature, etc.), measurement procedure, data processing, and the properties of the material. Different isotropic and anisotropic properties of these fabrics allow us to account for the effect of specular reflectance. Datasets B, C, D all consist of leaf reflectance of *Fagus sylvatica* measured on a mature tree over the growing season (B), on a French population (C), and on a set of Swiss populations (D). The *F. sylvatica* datasets incorporate different biological sources of variation as well as measurement uncertainty (shaded cells in table). Phenology, microclimate, herbivory and pathogens cause diverse leaf morphologies and chemical compositions between leaves of a same branch and between branches of a single individual. Genetic diversity between individuals creates an additional layer of biological variation among leaf traits. The spectral variation is approximated by the standard deviation and the coefficient of variation when normalized.

In a first step, we characterize the variation within an individual tree, considering a time series including spectral reflectance taken at different positions of the tree crown over the course of one growing season. Secondly, we disentangle the contribution of individual variation from the variation observed among forest sites, based on variation partitioning.

Lastly, we evaluate the influence of sample size on the observed variation using statistical approaches, and suggest measures towards a standardized protocol to allow rigorous quantification of biological variation via LOP.

## 2. Materials and Methods

### 2.1. Experimental design

The study combines four datasets (Fig. 1.a). The first dataset (dataset A) comprises reflectance of fabrics (see 2.2). The three other datasets comprise reflectance of *F. sylvatica* leaves that were collected on one individual across the growing season (dataset B), and on several individuals from a French forest (dataset C) or Swiss (dataset D) forest stands (see 2.3). Fabric and leaf reflectance were estimated from measurements performed with a field spectroradiometer coupled with either a leaf clip or an integrating sphere (Fig. 1.b; see 2.4-2.6). For each dataset, we calculated the spectral variation (see 2.7). In dataset A, we compare the spectral variation to the measurement uncertainty, while in the other datasets we partitioned the spectral variation into measurement uncertainty and biological variation. We further distinguish between the biological variation originating from different levels of biological organization (i.e., within- and between-individual variation) (Fig. 1.c-d)

### 2.2. Fabrics used as standard materials

We selected four different fabrics according to their optical properties (Fig S1):

1. A *Camouflage fabric* from SSZ Camouflage Technology AG: opaque, mostly isotropic reflectance, spectrum in the visible and in the near infrared ranges highly similar to vegetation.
2. A green translucent, woven, 100% *cotton fabric*, which reflects light isotropically.
3. A green opaque *plastic fabric* which reflects light with a specular component depending only on the zenith angle.
4. A green translucent *satin fabric* which reflects light anisotropically. It has a very strong specular component which depends both on zenith and azimuth angles.

The optical properties of each fabric are assumed to be the same everywhere on the piece of fabric and to not change over time. The materials were purchased at Alja Nouveau AG, Oerlikon, CH.

Reflectance of fabrics was estimated from measurements performed with both sampling devices, i.e., a leaf clip and an integrating sphere, which were successively coupled to the same ASD FieldSpec spectroradiometer (see 2.4. Instruments) to assess the measurement uncertainty associated with the material reflectance (**Fig. 1.b**).

### 2.3. Leaf samples

Leaves of an approximately 200-year-old beech tree located on the campus of the University of Zürich (47°23’44.7”N 8°32’58.1”E) were sampled weekly over the entire growing season, from the 3^rd^ of May to the 7^th^ of November 2018. Eight repeat sampling spots, comprising three shaded sampling spots located under the crown, and five sun-exposed sampling spots, were selected to assess the variation in LOP within an individual tree at each point in time, as well as over the season. Of the five sun-exposed sampling spots, three were located on the east side at 3, 6, and 12 m tree height, and two on the south side at 6 and 12 m tree height to account for potential effects of varied light exposition and sampling height on LOP. Each week (n=28), one twig was sampled at each sampling spot. Leaf reflectance of three leaves chosen one at the base, one at the middle, and one at the tip of each twig was acquired *in situ* using a leaf clip coupled with an ASD FieldSpec spectroradiometer (see 2.4. Instruments). *In-situ* measurements were used to characterize the biological variation within an individual tree. The leaves were stored and transported to the laboratory in dry ice protected from the light. Within three hours following the sampling, leaf reflectance was acquired once more in a dark laboratory simultaneously with a leaf clip and an integrating sphere, ensuring that there was no water film on the surface of the leaves prior to measurement. Laboratory measurements were compared to assess differences in leaf reflectance due to the use of different sampling devices. Because of the smaller size of the leaves at the beginning of the season, measurements in weeks 18 to 20 were carried out on three individual leaves belonging to the same node.

To assess the variation in LOP between individuals and between sites (Fig. 1.c-d), we collected leaves from 189 beech trees in la Massane in the French Pyrenees (42°28’28.0”N 3°01’11.7”E) (Dataset C) and from 75 beech trees located in five sites in Jura in northern Switzerland (47°31’9.3”N 7°37’55.5”E) (Dataset D) (Fig. S2). The sampling was conducted on 6-10 of July and on 25-26 of July 2019 for the French and Swiss forests, respectively.

From each sampled tree, we harvested one branch per tree from the top of canopy with a telescoping scythe (Takeni Trading co., Osaka, JP) in France and by helicopter in Switzerland, kindly allowed as part of the long-term monitoring sample harvest by the Institut für Angewandte Pflanzenbiologie (IAP). From each branch, we randomly selected three leaves from which we acquired leaf reflectance within three hours following the sampling using an ASD FieldSpec spectroradiometer (see 2.4 Instruments). Two ASD Fieldspec spectroradiometers were used, each measuring approx. half of all samples per site. One instrument was used for one plant individual.

### 2.4. Instruments

Reflectance was calculated from measurements performed with a FieldSpec spectroradiometer (ASD Inc., Boulder, CO, USA) coupled with a sampling device (leaf clip or integrating sphere). Datasets A and B were obtained using a FieldSpec 4 Wide-Res device (serial n°18130), while datasets C and D were measured using two FieldSpec 4 Standard-Res (serial n°18130 and 18140). A leaf clip, consisting of the plant probe plus leaf clip (model A122317, serial n°455 and 885, ASD Inc., Boulder, CO, USA), and/or an integrating sphere (serial n°6045-2, ASD Inc., Boulder, CO, USA) were used as sampling device depending on the sample and measurement procedure (see 2.5.

Measurement procedures). Both sampling devices have an internal standardized light source allowing the measurement to be taken independently of external illumination conditions. The light source of the leaf clip is a halogen bulb with color temperature of approximately 2900K, whereas the integrating sphere is supplied with a collimated tungsten light source (ASD Inc, 2008).

The FieldSpec spectroradiometer employs three detectors covering the visible and near infrared (VNIR) to the shortwave infrared (SWIR) range from 350 to 2500nm. The VNIR detector is a silicon photodiode array while the two SWIR detectors are thermo-electrically cooled indium gallium arsenide detectors, covering the 1001-1800 nm and 1801-2500 nm ranges, respectively. The detectors are covered by order separation filters and the light is dispersed by a holographic diffraction grating. The factory wavelength accuracy is 0.5 nm while the nominal spectral resolution ranges from 3 nm in the VNIR to 10 nm in the SWIR (ASD Inc, 2010).

In dataset A and B, optimization of the detector sensitivities using the standard light source and white background was set to 17 ms for the leaf clip and to 136 ms for the integrating sphere to maximize the detector sensitivity and minimize the exposure time.

Exposure time in case of biological material should be considered with care as prolonged exposure time to heat from the lamp, especially in the leaf clip set-up, may alter LOP. In this regard, optimized acquisition time was set to 8.5 ms in dataset C and D.

### 2.5. Measurement procedures

Measurements performed with a leaf clip consisted of four successive readings: the white reference (Rw), the white reference plus target (Tw), the black reference (Rb), and the black background reference plus target (Tb). Measurements performed with the integrating sphere followed the recommended method of the manufacturer (ASD Inc., 2008) consisting of three successive readings: the reflectance reference (Ir), the reflectance sample (Is) and the dark reading (Id). Uncalibrated Spectralon® panels (99% nominal reflectance) were used as a white reference for integrating sphere measurements.

Measurements of the fabrics were taken with both sampling devices in a dark laboratory. For these measurements, 15 scans per reading were recorded and averaged. The measurement was repeated on 6 different spots on the fabric sample. The sample was rotated after each measurement to account for the anisotropic properties of the material. One dark reading was performed per sample.

Measurements of leaf samples were taken *in situ* with a leaf clip as soon as possible after branches were retrieved from trees; in dataset B and C measurements were taken within 10 min and comprise 5 scans per readings, while in dataset D measurements were taken within 3 hours and comprise 5 scans per readings. Additional measurements of leaves of the individual tree (dataset B) were carried out in a dark laboratory using both sampling devices within three hours after sampling (leaves were kept on dry ice during these up to three hours; leaves were not frozen and no water film was present at the surface by the time of measurement). Measurements with a leaf clip and an integrating sphere were respectively taken on the right and left side of the leaf to avoid heating the leaf with repeated measurements on the same spot. For these measurements, 5 scans (10 scans, dataset D) per reading were recorded. The first and last scans were systematically removed before averaging the scans to avoid potential contamination from previous readings or from hasty opening of the sampling device.

Measurement results were saved as digital numbers (DN values), corresponding to the signal intensity after optimization, and as reflectance, corresponding to the signal intensity calibrated against the white reference by the ASD software. The reflectance obtained is a reflectance factor resulting from a bi-directional measurement (leaf clip) or a directional-hemispherical reflectance (integrating sphere) (c.f., Schaepman-Strub et al., 2006 for terminology). Reflectance units vary between 0 and 1, where 1 corresponds to 100% reflectance of the white reference of the sampling device.

### 2.6. Calculation of sample reflectance

The reflectance of a sample (R) is calculated from the mean of scans of the different readings (see 2.5 Measurement procedures). Several calculation methods are available in the literature using either the DN or the reflectance values (**Table S1**). To assess the systematic effects potentially induced by various calculation methods on the uncertainty associated with reflectance, we compared the reflectance of the fabric samples calculated from the equations shown in **Table S1**. The material reflectance values calculated from DN values and reflectances are identical when estimated from measurements with an integrating sphere and differ by up to 2.2% when estimated from measurements with a leaf clip **(Fig. S3)**.

The reflectance of all other samples was calculated from recorded reflectance values according to *equation 1* (leaf clip measurement) and *equation 2* (integrating sphere measurement). The (-) indicates that the result is dimensionless: a ratio between 0 and 1, referred to throughout this manuscript as reflectance units.

Leaf clip:

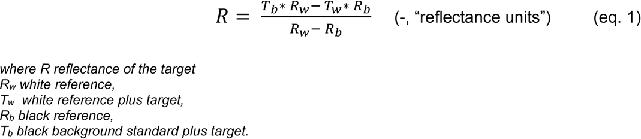

Integrating sphere:

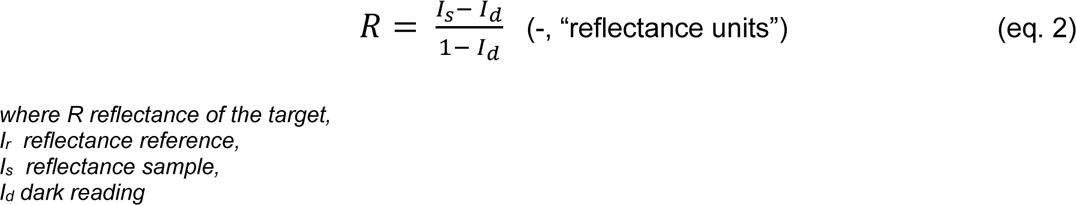

Radiometric steps commonly associated with incomplete warming-up of the instrument (ASD Inc, 2010) and ambient temperature (Hueni and Bialek, 2017) appeared at the detector limits (1000 and 1800 nm, respectively). We thus applied a corrective model developed by Hueni and Bialek (2017), originally targeting the correction of radiances. It was adapted within this study to also correct reflectance values where, previously, the correction values of the model were noise limited and thus introduced new artifacts (Hueni, 2021).

### 2.7. Calculation of measurement uncertainty, spectral, and (non-) biological variation

The measurement uncertainty, hereafter, corresponds to the combined uncertainty associated with the reflectance and calculated according to the law of propagation of uncertainties. The absolute measurement uncertainty (eq. 3) equals the standard uncertainties (Uxi) of the different readings involved in the reflectance calculation weighed by the corresponding coefficients of sensitivity (cxi) and added in quadrature. The standard uncertainty was defined as the standard deviation of the reading (STDxi) divided by the square root of the number of readings (N) (**Fig. S4**). The coefficient of sensitivity corresponds to the partial derivative of the reflectance with respect to the reading (xi) and describes how much the reflectance changes when the reading xi changes. Except for the two measurements including the target (i.e., the measured leaf or fabric), the uncertainties associated with each reading do not correlate with each other in any of the datasets (**Fig. S4-7**). Consequently, co-variation terms were neglected in the uncertainty calculation.

The relative measurement uncertainty was defined as the absolute uncertainty divided by the mean reflectance of the material (or across the dataset, for leaf measurements) with units of per cent (eq. 4).

Absolute measurement uncertainty:

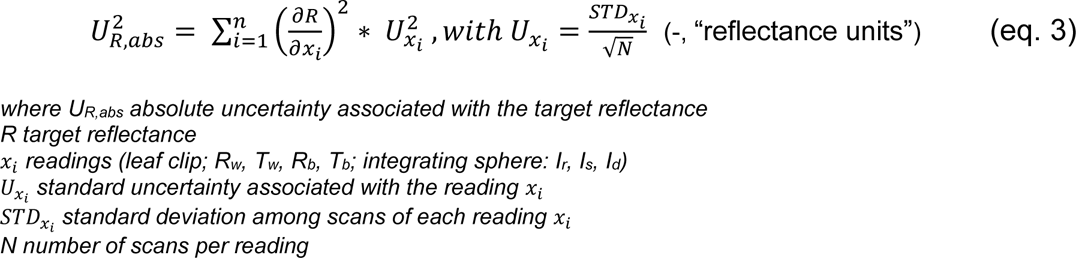

Relative measurement uncertainty:

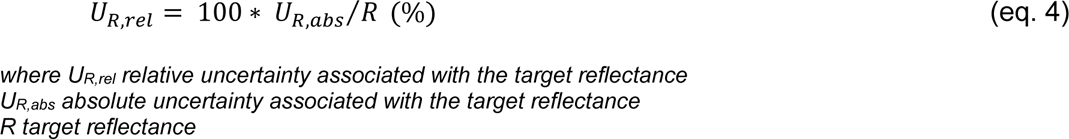

The standard uncertainty of each reading corresponds to the probability distributions associated with all different sources of uncertainty, including the instrument characteristics and experimental conditions. The contribution of individual sources of uncertainty was not considered in our uncertainty calculation, except for the ambient temperature (see 2.6).

Definitions and terminology are consistent with the Guide to the Expression of Uncertainty in Measurement (JCGM, 2008).

The measurement uncertainty was calculated for single and repeated measurements of the four fabrics (dataset A), successively recorded with a leaf clip and an integrating sphere. In the case of leaf datasets (dataset B, C, D), the measurement uncertainty was calculated for each single leaf measured. It was, however, not possible to establish the uncertainty associated with repeated measurements of leaves, as LOP vary from one individual leaf to another, and individual leaves were not measured repeatedly using the same set-up (except for the two measurements required to calculate reflectance). Additional measurements of the same leaf were conducted only in a limited manner, with extra precautions, to compare different sampling devices as described in sections 2.3-2.5. We also note that repeated measurements of the same leaf may incorporate stress responses of that leaf to the measurement, especially the strong light exposure, and thus could not be considered repeated measurements in the same way as repeated measurements of non-living fabrics. We can infer how measurement uncertainty may accumulate over many measurements from Dataset A, whereas we present mean and variance of calculated per-sample uncertainty for the leaf datasets.

The spectral variation was defined as the standard deviation among leaf reflectance within a dataset. The relative spectral variation, or coefficient of variation, corresponds to the standard deviation expressed as a percentage of the mean reflectance (eq. 5).

The (non-)biological variation was defined as the spectral variation from which we subtracted the mean uncertainty associated with a single measurement (eq. 6). Non-biological variation is here operationally defined as variation in optical properties of fabrics. We assumed that the optical properties of a standard fabric are homogenous. We thus expect the non-biological variation to be nil or almost nil, and assess this by comparing the deviation of fabric measurements against their calculated measurement uncertainty.

Biological variation is attributed to variation in LOP caused by differences in leaf morphology and chemistry.

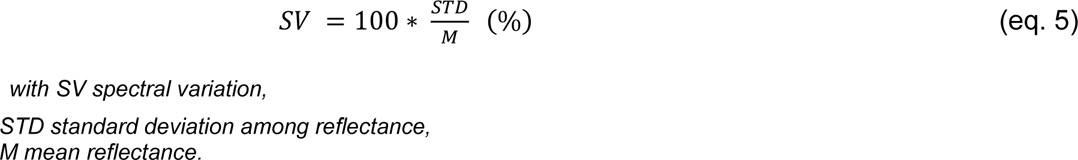

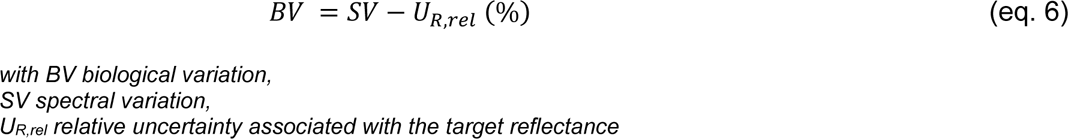

### 2.8. Data treatment & Statistical analyses

Data represent mean values ± standard deviation. The 95% confidence interval corresponds to two standard deviations under the normality assumption. The number of replicates is indicated for each dataset. Data processing and statistical analyses were all performed in Matlab R2020a. Normality tests, t-tests and ANOVAs were performed on individual wavelengths; see Results and figure captions for details. Scripts and source data are available in Dryad (https://doi.org/10.5061/dryad.gtht76hkx). FieldSpec spectroradiometer data are also deposited in SPECCHIO (http://sc22.geo.uzh.ch:8080/SPECCHIO_Web_Interface/search, Hueni et al., 2020) and can be found with the identifiers ‘Field spectroscopy Fabrics’ (dataset A), ‘Field spectroscopy F. Sylvatica individual’ (dataset B), ‘Field spectroscopy F. sylvatica La Massane’ (dataset C), ‘Field spectroscopy F. sylvatica SwissForest’ (dataset D). Visualization and descriptive statistics were performed with Originlab 2020.

## 3. Results

### 3.1. Uncertainty and variation associated with measurements of fabric standards

We first characterized the uncertainty associated with the measurement of fabrics.

Unlike LOP, which are known to be heterogenous within a leaf and sensitive to light exposure time (Jacquemoud and Ustin, 2019), optical properties of fabrics are assumed to be homogenous and constant over time. Also, the measurement of these fabrics under controlled laboratory conditions allows us to isolate the spectral variation which can be attributed to the measurement uncertainty related to the spectroradiometer and the sampling device. We explore two levels of uncertainty: the uncertainty associated with a single measurement, and the uncertainty generated by repeated measurements of the same fabric. Both measurement uncertainties are combined uncertainties calculated on four fabrics successively measured with a leaf clip, and an integrating sphere.

#### 3.1.1. Measurement uncertainty associated with the leaf clip

The measurement uncertainty associated with a single leaf clip measurement of the camouflage, plastic, and cotton fabrics represented less than 0.002 reflectance units across the full spectral range (**Fig. 2e-g**). In contrast, the satin fabric (**Fig. 2h**) had a high specular component, and its uncertainty averaged at 0.1 reflectance units in the SWIR. The uncertainty associated with a single measurement of the material reflectance was also of the same magnitude as the standard uncertainty (i.e., standard deviation) associated with the white reference (0.0013 ± 0.0003) (**Fig. S4**), except for the anisotropic fabric (satin).

**Figure 2.**
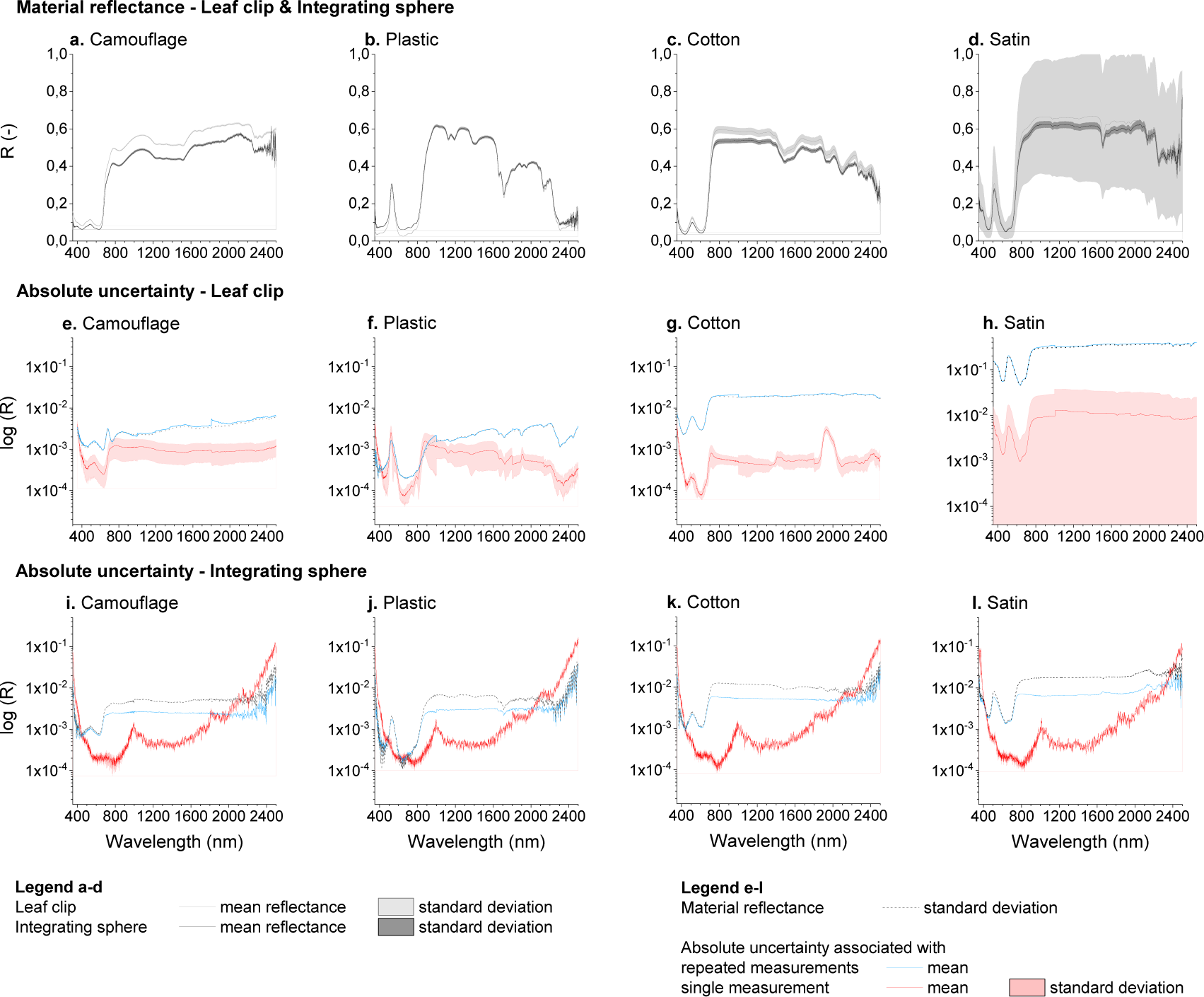
Spectral variation and measurement uncertainty associated with the reflectance of four fabrics. The mean reflectance and standard deviation of four fabrics (camouflage material (a), plastic material (b), cotton fabric (c), satin fabric (d)) measured with a leaf clip *(grey)* and an integrating sphere *(dark)* indicate the spectral variation obtained by using different sampling devices. The spectral variation, or standard deviation (*dotted line*), obtained for each combination of fabric and sampling device is compared to the corresponding measurement uncertainties associated with single and repeated measurements (e-l). The absolute uncertainty calculated for each single measurement according to the law of propagation of errors is shown as the mean (n=6) of these calculations *(red)* and their standard deviation *(red shade)*. The absolute measurement uncertainty associated with repeated measurements *(blue)* of each of the four standard materials measured with a leaf clip (e-h) and an integrating sphere (i-l) includes all measurements taken for each material (no standard deviation).

Repeated measurements of the same fabric increased the measurement uncertainty such that it became nearly identical to the standard deviation of measurement, which was lowest in the VNIR and ranged from about 0.0001 to 0.01 reflectance units for all but the satin fabric. The uncertainty increased with increasing wavelength in the SWIR for all materials. The leaf clip allows for a bi-directional measurement. As the sample was rotated in the leaf clip after each measurement, the high zenith anisotropy of the satin caused a high standard deviation. Thus, the measurement uncertainty associated with the fabric measurements appeared to strongly depend on the optical properties of the fabric itself, and overall to be similar to the standard deviation from repeated measurements of the same fabric.

#### 3.1.2. Measurement uncertainty associated with the integrating sphere

The measurement uncertainty associated with a single integrating sphere measurement was similar for all four standard materials (**Fig. 2i-l**). It comprised between 0.0001 and 0.001 reflectance units in the spectral range of 400-2000 nm and abruptly increased up to 0.1 reflectance units at both ends of the spectral measuring range (i.e., 350-400 nm and 2000-2500 nm). Irrespective of the fabric measured, the measurement uncertainty associated with repeated measurements was between 0.001 and 0.01 reflectance units, slightly less than the standard deviation. The measurement uncertainty was systematically lower in the VNIR (0.001 ± 0.0005 reflectance units) than in the SWIR range (0.004 ± 0.002 reflectance units). Repeated measurements also helped to reduce the measurement uncertainty at both ends of the spectral measuring range by an order of magnitude compared to a single measurement. These results indicate that the anisotropic properties of the fabric did not affect the measurement uncertainty associated with the integrating sphere measurements, likely because the integrating sphere integrates the reflected light over a full hemisphere. In other words, the measurement uncertainty seemed to depend more on the measurement conditions and the characteristics of the spectroradiometer and the integrating sphere, than on the optical properties of the fabric.

#### 3.1.3. Measurement uncertainty and non-biological variation when using both sampling devices

The measurement uncertainty associated with single measurements of all fabrics using either a leaf clip or an integrating sphere ranges from 10^-4^ to 10^-1^ reflectance units and tends to be independent of the specular component (except for satin). Repeated measurements of the same fabric were generally associated with a greater uncertainty than single measurements. Differences between uncertainties associated with single and repeated measurements tend to increase for fabrics with a stronger specular component, though to a lesser extent for measurements using an integrating sphere compared to those using a leaf clip. The reflectance of opaque and more isotropic materials (camouflage and plastic) calculated from measurements performed with either a leaf clip or an integrating sphere led to comparable measurement uncertainties. However, only measurements performed with an integrating sphere ensure a low measurement uncertainty (< 0.01 reflectance units) associated with the reflectance of translucent and strongly anisotropic materials (cotton and satin). The integrating sphere thus allows for more comparable results between materials than the leaf clip, although it should be noted that there are applications in which the anisotropic reflectance captured by the leaf clip and not by the integrating sphere may be important information. Besides, there is a systematically higher mean reflectance obtained with the leaf clip compared to the integrating sphere (**Fig. 2a-d**). Systematic differences in reflectance and magnitude of uncertainty associated with measurements performed using either a leaf clip or an integrating sphere indicate that the magnitude and distribution of measurement uncertainty is sampling-device specific.

The uncertainty associated with repeated measurements using either sampling device was similar to the standard deviation. The non-biological variation corresponding to the difference between the standard deviation and the measurement uncertainty was nil or almost nil, which supports the assumption that the optical properties of a fabric are homogenous. Also, while the uncertainty associated with a single measurement mainly accounts for the variation due to the used instrumentation, the uncertainty associated with repeated measurements accounts for the variation due to measurement conditions and the specular component of the target material. Thus, the standard deviation appears to be an adequate proxy for spectral variation that includes the non-biological variation (here, nil or almost nil) and the measurement uncertainty, encompassing the variation due to the instrumentation, the measurement conditions, and the anisotropic directional reflectance behavior (see **Fig. 1c**).

### 3.2. Uncertainty and variation associated with measurement of leaves

We inferred the biological variation from the spectral variation in leaf reflectance (see **Fig. 1c**), hereafter approximated by the coefficient of variation (CV). As the accuracy of the biological variation inferred from leaf reflectance depends on our ability to isolate the measurement uncertainty from the spectral variation, we quantified the contribution of the measurement uncertainty to the spectral variation observed in our three datasets, i.e., *F. sylvatica* individual (over time, B), French forest (single site, C), and Swiss forest (multiple sites, D). As the heat of the light source and the leaf holder may damage the leaf over time, only single measurements comprising repeated scans are generally performed. Also, we restricted the calculation of the measurement uncertainty associated with leaf reflectance to the uncertainty associated with a single measurement, defined as the mean of multiple scans per leaf as described in section 2.7. Uncertainty calculation on fabrics (see 3.1) showed that the uncertainty associated to a single measurement accounts for the variation induced by the instrumentation and measurement conditions, but underestimates the uncertainty associated with repeated measurements that depends on the specular component of the target. Thus, the resulting spectral variation encompassed biological variation and potential anisotropic directional reflectance behaviour of the leaf. Nevertheless, given that the measurement uncertainty associated with single measurement of leaf reflectance was comparable to isotropic standard materials, we neglected the contribution of the specular component and defined the biological variation as the spectral variation from which the mean uncertainty associated with single measurements was subtracted. We further investigated the contribution of various levels of biological organisation within and between individuals to the observed biological variation (see 3.2.3).

#### 3.2.1. Measurement uncertainty associated with leaf reflectance

In dataset B, we successively measured the leaves with a leaf clip and an integrating sphere, taking specific precautions against leaf degradation as described in sections 2.3-2.5, and characterized the associated measurement uncertainties (**Fig. 3**).

**Figure 3.**
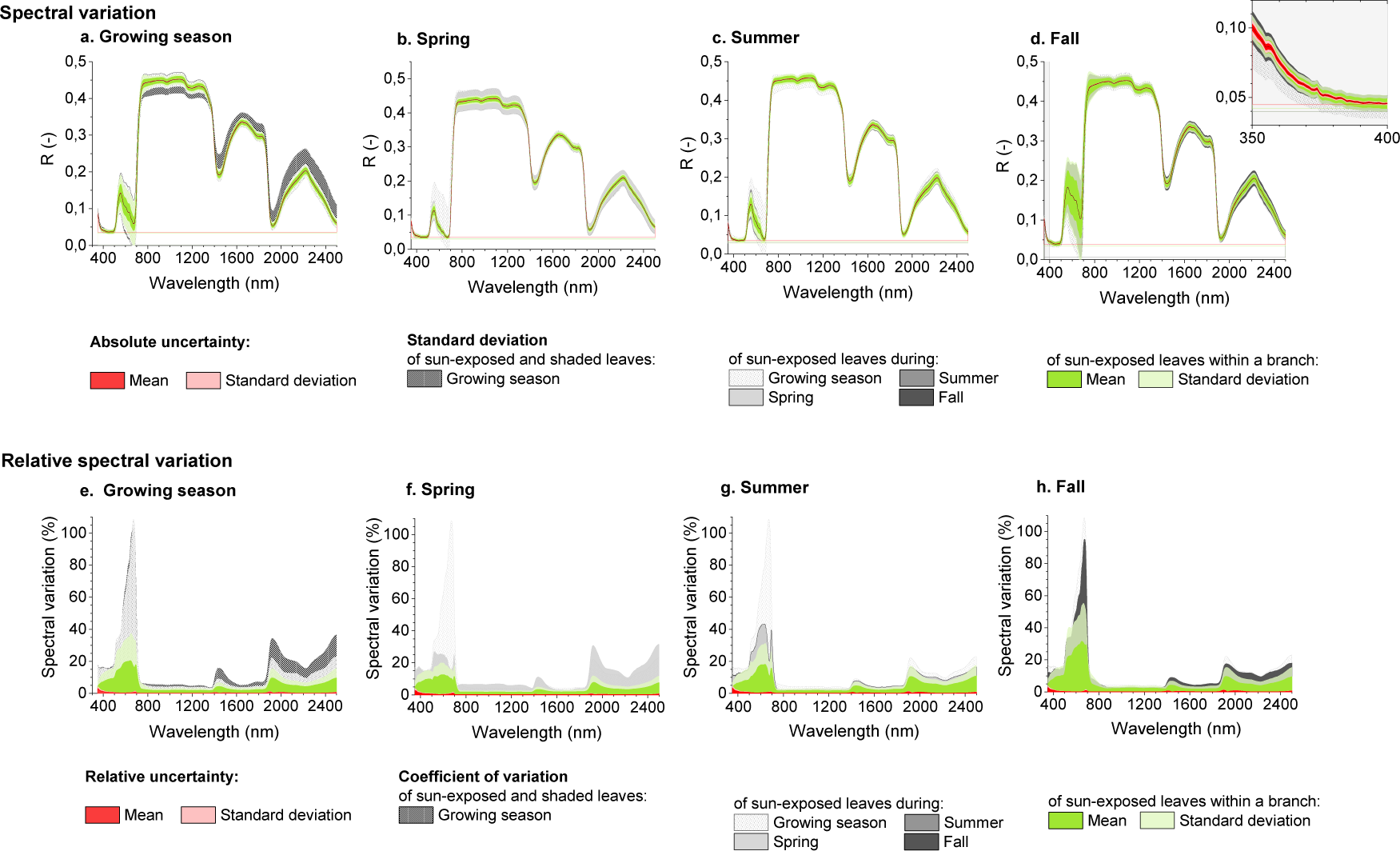
Spectral variation and measurement uncertainties of shaded and sun-exposed leaves of a *Fagus sylvatica* individual in spring, summer, and fall. The first row represents the mean leaf reflectance *(black)* of shaded and sun-exposed leaves of a *Fagus sylvatica* individual across the growing season (a), and more specifically of sun-exposed leaves in spring (b), summer (c), and fall (d). The red shades represent the mean measurement uncertainty associated with leaf reflectance and its standard deviation. The dark and light green shades represent the mean spectral variation within a branch composed of 3 leaves and the corresponding standard deviation. The dotted shade represents the spectral variation as calculated by the standard deviation observed among sun-exposed and shaded leaves over the growing season, while the light and dark grey shades represent the spectral variation observed among sun-exposed leaves (n= 5 branches) either across the growing season or at a specific phenological phase. The second row represents the relative uncertainty *(red)* and the coefficient of variation associated with the variation within a branch *(green)* and within the tree *(grey)* across the season (e), in spring (f), summer (g), and fall (h).

The uncertainty associated with the leaf clip measurements averaged at 0.0004 ± 0.0002 reflectance units (**Fig. 3a**), corresponding to 0.3% of the leaf reflectance on average across the full spectral range. The measurement uncertainty varied between 0.2 and 3% of the mean reflectance in the VNIR and averaged at 0.08% in the SWIR (**Fig. 3e**). The absolute uncertainty when measuring leaves with an integrating sphere was statistically identical to that of standard materials (ANOVA, df=26, p>0.5), confirming that the measurement uncertainty for integrating sphere measurements was independent from the target.

In contrast, the difference between leaf clip and integrating sphere measurements represented up to 40-80% of the mean leaf reflectance in the VNIR and between 10-40% of the mean reflectance in the SWIR range, which was twice as large as the relative difference observed on standard materials, confirming that the relationship between data obtained with these two different sampling devices is target-specific (**Fig. S10**). The largest differences were observed at wavelengths generally used to retrieve pigment and water contents.

In the following sections, we investigate relationships between measurement uncertainty and variation (spectral and biological) among leaf reflectance calculated from leaf clip measurements only.

#### 3.2.2. Measurement uncertainty and biological variation within an individual

We investigated dataset B, comprising leaf reflectance spectra from a *F. sylvatica* individual sampled across one growing season. We calculated the spectral variation and characterized the measurement uncertainty and biological variation associated with leaf reflectance.

##### 3.2.2.1. Spectral variation and measurement uncertainty associated with leaf reflectance within an individual

We calculated the spectral variation among leaves of an *F. sylvatica* individual sampled weekly during the growing season, including sun-exposed and shaded leaves (**Fig. 3a**). In addition, we considered the spectral variation among sun-exposed leaves only, divided into measurements conducted in spring, summer and fall (**Fig. 3b-d**). The largest contribution of the measurement uncertainty to the spectral variation (20%) was observed for spectral bands below 400 nm where the instrument noise is governed by a low quantum efficiency of the silicon VNIR detector combined with a low intensity of the halogen lamp used as light source. Above 400 nm, the relative uncertainty represented on average 2.5 ± 1.6% of the spectral variation among sun-exposed leaves (**Fig. 3e**). Although the measurement uncertainty tended to contribute more to the spectral variation in summer (3.5 ± 1.8%) and fall (3.0 ± 1.8%) than in spring (1.9 ± 1.0%), its contribution to the spectral variation was statistically independent of the time of measurement (ANOVA, df=198, p>0.5) across the full spectral range (**Fig. 3f-h**). Similarly, the relative uncertainty represented on average 5.6 ± 3.1% of the spectral variation among sun-exposed leaves belonging to the same branch. Maxima in measurement uncertainty roughly corresponded to maxima in spectral variation (e.g., at 690, 1415, 1900 nm) and did not exceed 10% and 20% of the spectral variation observed within an individual and within branches, respectively.

##### 3.2.2.2. Spectral variation and biological variation within an individual

We defined the biological variation among leaves of the *F. sylvatica* individual as the relative spectral variation calculated as coefficient of variation from which we subtracted the relative measurement uncertainty. **Fig. 4a** illustrates how the biological variation among sun-exposed and shaded leaves evolved across weeks during the growing season. The biological variation in the VNIR region varied from as low as 5% to as high as 110% with respect to the wavelength and the sampling time. Biological variation was highest in the spectral range of 500-750 nm corresponding to pigment absorption, and showed a clear time dependency. We distinguished three time periods, roughly following the main phenological stages of a leaf, i.e., the leaf development in spring (week 1-9), the maturation of fully developed leaves in summer (week 10-19), and leaf senescence in fall (week 20-28). Across the spectrum, the mean biological variation was 20% of the leaf reflectance in spring, between 30% and 50% in summer, and over 100% in fall, revealing an increasing diversity in spectral features within a tree as leaves mature and senesce. However, the biological variation at a given wavelength remains largely constant (< ± 10%) over time in the SWIR region, and thus the time dependency is driven by differences in the VNIR. The largest values in the SWIR (15%-40%), mainly driven by the water content, appeared between 1400-1500 nm and between 1800-2500 nm. Out of a biological variation of 40% in the SWIR, up to 30% originate from the differences between sun-exposed and shaded leaves (**Fig. 4b-c**).

**Figure 4.**
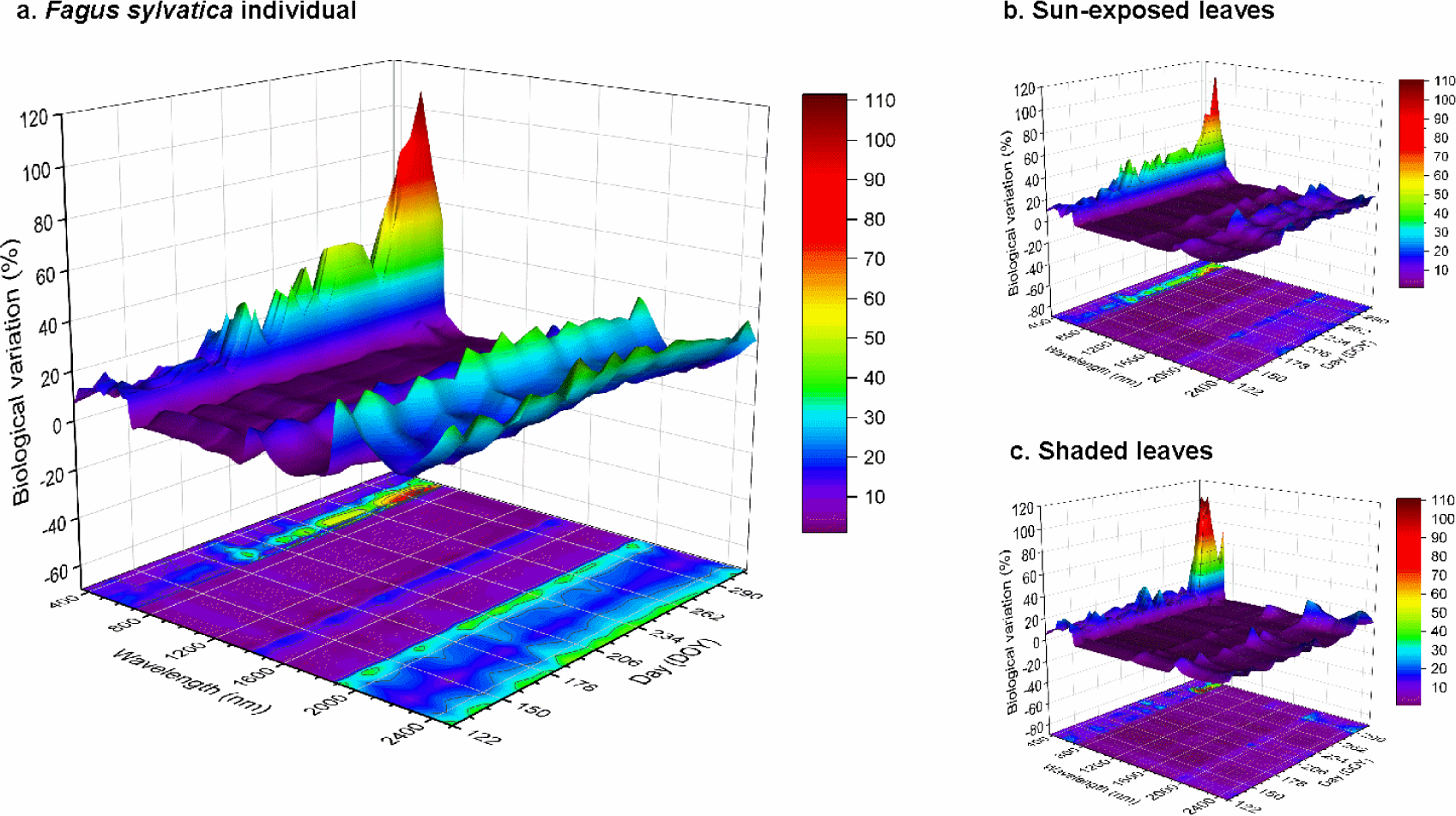
Biological variation within a *Fagus sylvatica* individual during the growing season retrieved from leaf reflectance. The biological variation was obtained by subtracting the measurement uncertainty from the spectral variation and describes the spectral diversity among weekly sampled (a) sun-exposed and shaded leaves (n=24), (b) sun-exposed leaves (n=15), (c) and shaded leaves (n=9). The biological variation of individual sampling spots is available in the supplementary data (**Fig. S11**).

Among individual sun-exposed branches only, the biological variation across the growing season comprised between 1.5%-10% in the SWIR, and averaged at 11± 6% in the VNIR region (**Fig. S11**). The biological variation in the VNIR was minimal in spring (9±3%) and had its maximum in fall (16±10%). We observed the same pattern above 1800 nm in the SWIR, though smaller in magnitude than in the VNIR. The biological variation within a branch explained on average 40±6% of the biological variation observed within a tree (**Fig. 3e-h**). Its contribution was minimal in spring (32±12%) and maximal in autumn (54±16%). The remaining biological variation (2-100%) originated from differences in leaf reflectance between sun-exposed branches.

#### 3.2.3. Measurement uncertainty and biological variation among several individuals

We characterized the spectral variation, measurement uncertainty and biological variation in datasets C and D, which consist of *F. sylvatica* trees sampled within a week in a French forest (C) and at 5 Swiss forest sites (D). We further partition the biological variation into variation within tree, among trees and among sites.

##### 3.2.3.1. Spectral variation and measurement uncertainty associated with leaf reflectance among several individuals

In both datasets C and D, the spectral variation defined as the coefficient of variation represented between 4% - 35% of the mean reflectance (**Fig. 5**). Similar to dataset B (single individual), the spectral variation was largest between 400 and 700 nm in the VNIR, and centered at 1145, 1930 and beyond 2400 nm in the SWIR.

**Figure 5.**
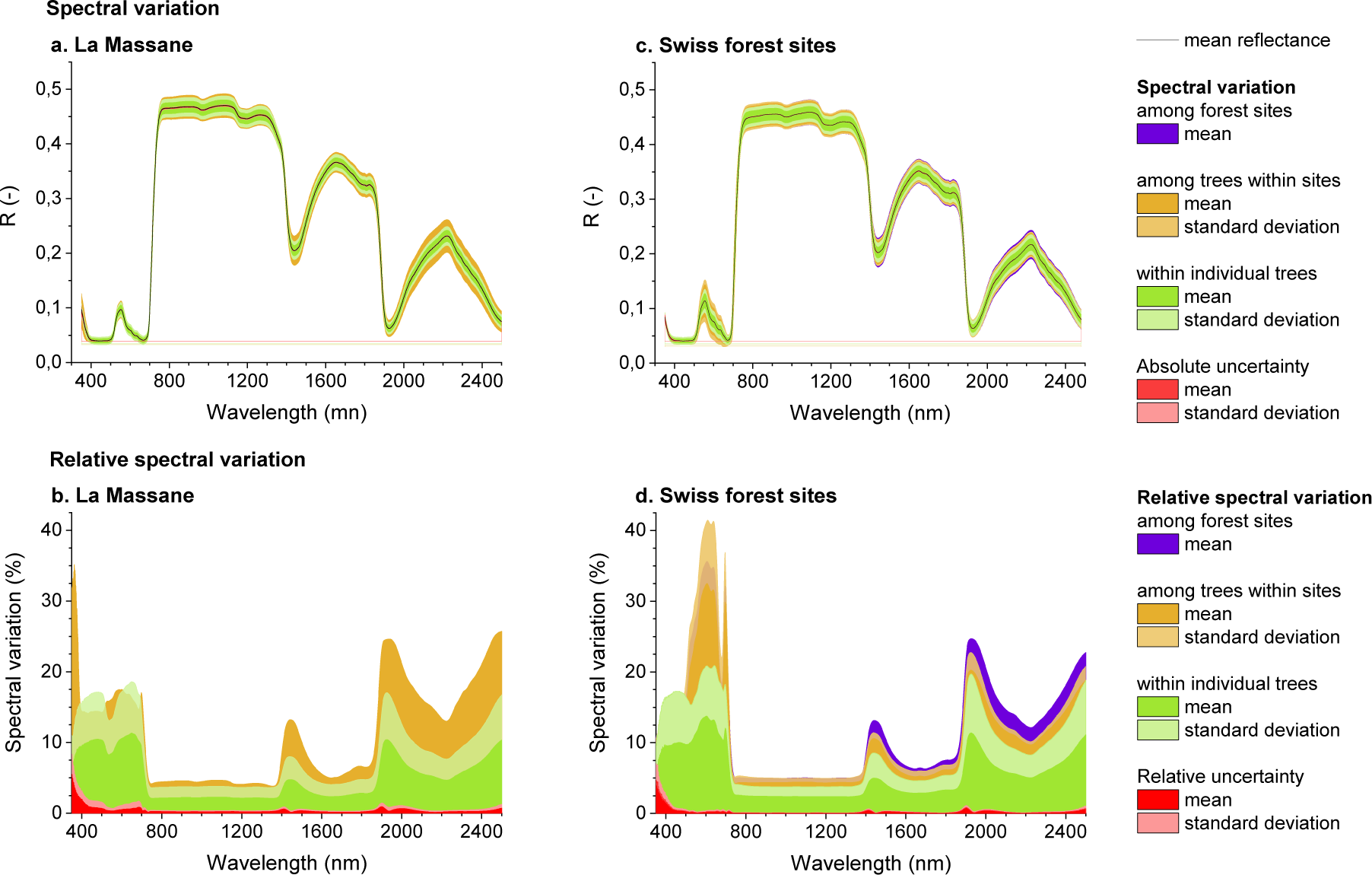
Spectral variation and measurement uncertainty within trees, among trees, and among sites for *Fagus sylvatica* leaves measured in a French (a-b) and five Swiss forest sites (c-d). On the first row the black line represents the mean reflectance of each dataset, while the red areas represent the mean measurement uncertainty and corresponding standard deviation. The green, orange and purple areas represent the mean (*opaque area*) and standard deviation (*transparent area*) of the spectral variation (calculated as standard deviation) within a tree, between trees within a site and among sites, respectively. The second row represents the relative uncertainty *(red)* and the coefficient of variation associated with the variation within a branch *(green)* and within the tree *(grey)* across the season (e), in spring (f), summer (g), and fall (h). The 95% confidence interval was plotted independently in the supplementary data (**Fig. S12**).

The maximum uncertainty was observed below 400 nm (French site: 5.5%, Swiss sites: 4.7%). After subtracting the measurement uncertainty, the spectral variation below 400 nm still exceeded 30% of the mean reflectance. This variation is unlikely to be due to biological variation, but rather due to other factors. We expect a lower signal-to-noise ratio near the limit of detection and the atmospheric scattering of light in blue wavelengths that may enter around the edge of the leaf clip. Consequently, spectral variation below 400 nm was excluded from the following analysis.

The relative uncertainty above 400 nm in datasets C and D was on average equal to 0.3 ± 0.2% and 0.2 ± 0.2% of reflectance, representing on average 3.4 ± 1.7% and 1.5 ± 1.3% of the spectral variation, respectively. The relative uncertainty represents up to 13% (dataset C and D) of the spectral variation in the VNIR, and up to 7.8% (at 1380nm, dataset C) and 4.5% (at 1400nm, dataset D) in the SWIR.

##### 3.2.3.2. Spectral variation and biological variation among several individuals

Within the spectral variation associated with biological variation (after subtracting the estimated measurement uncertainty), we differentiated three levels of biological organisation: (i) the individual level, representing the biological variation between leaves belonging to a single tree; (ii) the single-site level, corresponding the biological variation between trees belonging to a single forest site; and (iii) the multiple-site level, comparing the variation observed between trees belonging to several forest sites (in dataset D only). The single- and multiple-site comparisons evenly incorporate variation due to the parallel use of two radiospectrometers, whereas the within-individual variation for each site may be overestimated as a result of using measurements from two instruments (all leaves for one individual were randomly measured with either one or the other instrument). The spectral variation calculated for a single instrument on an individual sampling day was comparable to the spectral variation observed in the individual (B) and Swiss forest site (D) datasets (**Fig. S8**).

Maxima of biological variation in the VNIR tended to differ between levels of biological organization and were located at 480 and 660 nm for the individual level, and at 608 and 695 nm for the single- and multiple-site level. In the SWIR, maxima of biological variation were consistently observed across all levels of biological organization at 1451, 1981, and 2500 nm (**Fig. 5**). Interestingly, the biological variation within individuals was statistically identical between all datasets (B, C, D; ANOVA, df=223, p>0.05) across the full spectral range.

Biological variation within individuals ranged from 2 to 14% of the mean reflectance and mainly differed among datasets in the visible range of the spectrum. Within-individual variation represented 46 ± 10% of the spectral variation on average across the full spectral range.

The relative spectral variation between trees within a single Swiss forest site comprised between 12%-32% (versus 14-18% at the French site) of the mean reflectance in the visible range and between 4%-20% (4-25%, French site) of the mean reflectance in the NIR and SWIR (**Fig. 5**). After subtraction of the within-individual variation and measurement uncertainty, variation between trees belonging to a single site represented between 30-50% of the remaining spectral variation in the SWIR.

The relative spectral variation between trees growing in different Swiss forest sites represented up to 16% of the mean reflectance in the visible range and 22% of the mean reflectance in the SWIR, which corresponds to 15 ± 5% of the spectral variation on average. When considering biological variation within individual trees and sites, only the variation in the SWIR could be used to differentiate trees belonging to different sites of the Swiss forest dataset (dataset D).

##### 3.2.3.3. Systematic effect: sample size considerations

As observed in the Swiss forest dataset (dataset D), the biological variation observed at lower levels of biological organization may affect conclusions at higher levels. The quality of the information thus depends on our ability to accurately estimate mean biological variation and its standard deviation. We assessed the effect of sample size on the mean and standard deviation estimates of the spectral variation in the French forest (dataset C) consisting of 180 trees (**Fig. 6**).

**Figure 6.**
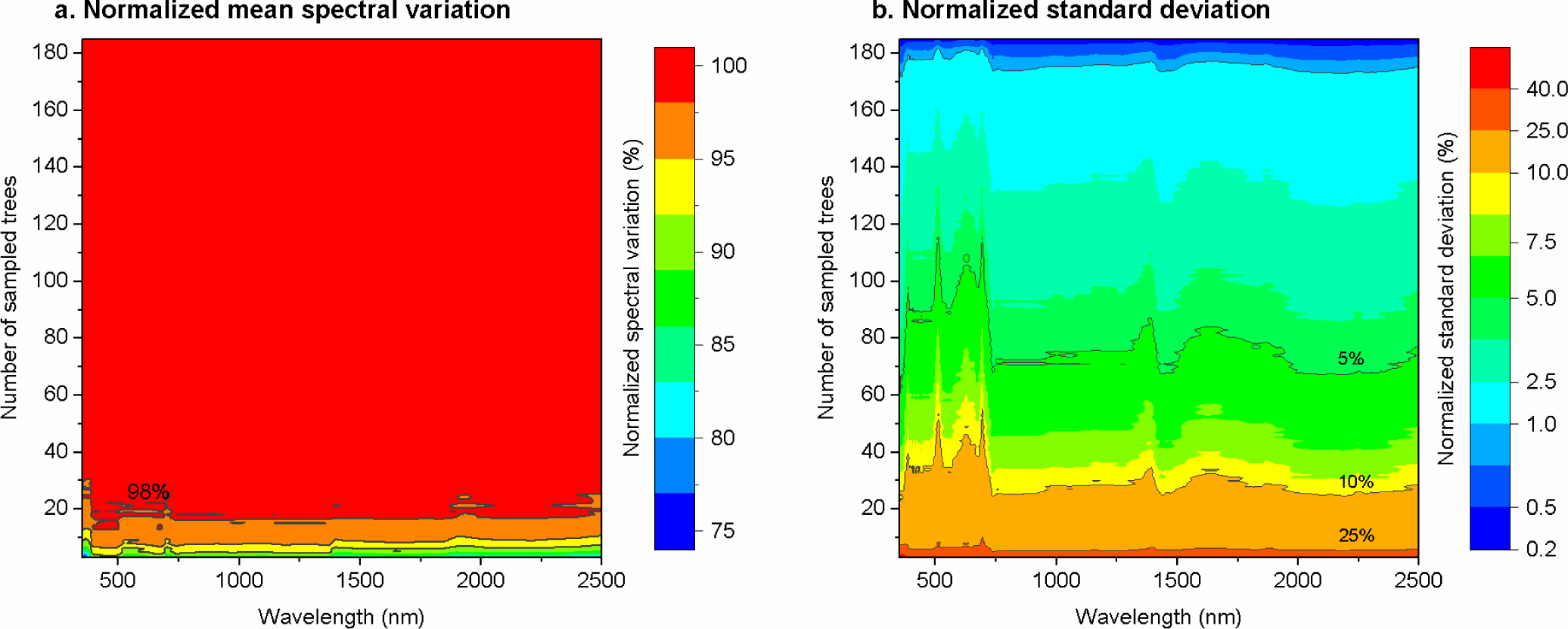
Effect of sample size on the mean spectral variation estimate and its standard deviation. The mean spectral variation (a) and its standard deviation (b) obtained for each sample size was normalized by the spectral variation and standard deviation observed in dataset C (n=180 trees) and by the mean reflectance. By increasing the number of trees sampled, the simulated mean spectral variation increased toward the observed mean variation while the simulated standard deviation decreased toward the standard deviation calculated on the full sample by bootstrapping.

The accumulation curve suggests that a minimum of 20 trees is required to approximate the mean biological variation originally observed among 180 trees of the French forest (dataset C) at a precision of ± 2%. However, in the SWIR, 70-90 trees and in the VNIR, 90-120 trees are required to obtain a standard deviation equal at ± 5% to the standard deviation calculated by bootstrapping over the entire sampled population of 180 trees.

## 4. Discussion

### 4.1. Sources of uncertainty in optical properties measurements

Measurement uncertainty is inherent to any optical measurement and characterizes the dispersion in reflectance that could reasonably be attributed to the target (JCGM, 2008). The optical properties of the target itself, as well as external factors, hereafter referred as sources of uncertainty, may impact the dispersion in reflectance. The spectral variation of the white background of the leaf clip reveals three sources of uncertainty generally affecting optical measurements (**Fig. S8**).

A first source of uncertainty is related to the optical sensor. ASD field spectroradiometers employ three detectors covering (i) the visible and near infrared (VNIR) from 350 to 1000 nm, (ii) the shortwave infrared from 1001 to 1800 nm (SWIR1), and (iii) from 1801 to 2500 nm (SWIR2). The largest variation (and thus uncertainty) occurs at the detector limits where the signal-to-noise ratio is the lowest (lowest DN values). Signal-to-noise ratio is a combination of instrument radiometric sensitivity and the intensity of the light source (Schaepman and Dangel, 2000). This intensity is low at the start of the VNIR and at the end of SWIR2 detector, following in general the Planck law. Unlike the second and third detector, the first detector (VNIR) is not cooled (ASD Inc., 2010). The sensitivity of the silicon of the first detector to temperature variation may in addition explain a higher spectral variation, particularly at the lower detector wavelength limit (Hueni and Bialek, 2017; Schaepman and Dangel, 2000). Variation in reflectance of the white reference, potentially altered by the instability of the detector over time (3 hours) was usually less than ±2% within one measurement session (**Fig. S9**).

A second source of uncertainty was associated with experimental conditions including moisture and ambient temperature. Although the internal calibration of the ASD spectroradiometer from DN values to radiance units eliminated most of the variation, post-correction based on temperature effect modelling was required to correct for the radiometric steps between the three individual detectors of the ASD full range instruments at 1000 nm and 1800 nm, respectively (Hueni and Bialek, 2017). While post-measurement corrections of known systematic effects help to remove biases in the measurement, they may also add uncertainties due to the underlying correction model, and should be applied with caution.

A third and related source of uncertainty is the potential contamination of the sampling device with previous samples, external light pollution, changing temperatures and atmospheric moisture content over time. Besides, multiple spectroradiometers may be used in parallel in order to achieve sufficient throughput during field sampling. These factors likely explain the larger variation in datasets B, C, and D, compared to measurements made in dataset A under laboratory conditions. The potential contamination of the white and the black references of the leaf clip over time with leaf material when measuring leaves is incorporated into uncertainty calculations using these references. In contrast, uncertainty calculations do not account for spectral variation potentially generated by several operators using multiple spectroradiometers. The French dataset (C) collected over three sampling days by two operators using two spectroradiometers in parallel show the largest spectral variation (up to 20%) in the spectral range 350-400nm where the signal-to-noise ratio is the lowest. The spectral variation calculated for a single instrument on an individual sampling day was comparable to the spectral variation observed in the individual (B) and Swiss forest site (D) datasets (**Fig S8**). In other words, the spectral variation in the French dataset reveals the systematic error introduced by two operators using two spectroradiometers in parallel (e.g., calibration, optimization, warming-up, detector characteristics, etc.) in changing measurement conditions (e.g., temperature, moisture, etc.). Accordingly, Lukěs et al. (2017) and Kuester et al. (2001) reported relative differences between ASD FieldSpec spectroradiometers of 1% to 2%, respectively, while Helder et al. (2012) estimated that the uncertainty induced by different operators was equivalent to 1.5% to 2.5% when best measurement practices are followed (cf., Hueni et al., 2017, for best practices framework).

Among all datasets, the VNIR spectral region appears to be the most sensitive to changes in measurement conditions. Nevertheless, measuring in reflectance mode, which requires initial normalization to the white reference and usually also entails periodic re-normalization every several measurements, corrects for most of the sources of uncertainty mentioned above and results in a spectral variation of the white reference below 1% in the VNIR (>400 nm) and 0.6% in the SWIR region (**Fig. 2**). The high variation remaining under 400 nm appears to be associated with a low signal-to-noise ratio of the detector, especially for one of our field spectroradiometers.

### 4.2. Measurement uncertainty on standard and leaf materials

Measurement uncertainty associated with optical properties appeared to not only depend on the characteristics of the spectroradiometer and its sensitivity to the measurement conditions, but also on the characteristics of the sampling device (leaf clip or integrating sphere). Uncertainty associated with leaf clip measurements additionally depended on the optical properties of the target material. Due to its geometry, a leaf clip device characterizes a bi-directional reflectance sensitive to the anisotropic properties of the target material; a stronger anisotropic directional reflectance behavior (specular component) results in a larger dispersion in reflectance (Schaepman-Strub et al., 2006). The hemispheric geometry of the integrating sphere offers a great alternative to the leaf clip device, allowing for measurements free of anisotropic directional reflectance behavior (Milton et al., 2009). Nevertheless, we showed that repeated measurements of the same target material rotated in the leaf clip device after each measurement reduced the directional effect and enabled us to account for the contribution of the specular component to the spectral variation and thus better inform us on the optical properties of the target material.

While some sources of uncertainty are well known and easily recognizable, assessing their individual contribution to the overall uncertainty budget can remain challenging due to their interdependencies (Kuester et al., 2001). We approached the uncertainty budget practically as an overall envelope calculated directly from measurements, rather than a detailed budget, thus providing a fast and reliable method to calculate the overall uncertainty associated with spectral measurements.

The measurement uncertainty associated with single measurements of all standard fabrics using either a leaf clip or an integrating sphere was very similar (except satin due to its specular qualities), suggesting that our calculation method is robust, suitable to assess uncertainty caused by instrumentation properties, and transferable to leaf measurements.

Repeated measurements of the same material were generally associated with a larger uncertainty than single measurements as the uncertainty accounts for the variation introduced by (even small) anisotropic properties of the target. To assess the contribution of the specular component to the spectral variation, we recommend repeated measurements with a leaf clip device. However, it must be noted that prolonged exposure to the light beam may alter leaf chemical composition (e.g., pigment degradation, water content).

The details of the approach we adopted to calculate the uncertainty budget are specific to the spectroradiometer-sampling device-target combination and would consequently change if either the spectroradiometer, the sampling device or the target change, as is apparent in the several combinations used in this study. Similarly, a systematic bias of the mean reflectance of an identical target was also observed by several authors when different spectroradiometers, sampling devices or protocols were used (Hovi et al., 2017; Pimstein et al., 2011; Jung et al., 2009; Castro-Esau et al., 2005). However, systematic differences due to the use of different spectroradiometers are often considered negligible (Lukes et al., 2017). Still, all of these biases create challenges for comparing data from different datasets and spectral databases. In a meta-analysis, Meireles et al., (2020) successfully harmonized leaf spectra measured using spectroradiometers from different manufacturers by applying a partial least square model. Errors between predicted and measured spectra remained below 0.01 (VNIR) and 0.03 (SWIR) reflectance units, corresponding to a spectral variation of up to 7-28% in the VNIR and 2-55% in the SWIR spectral range. It is worth noting that such variation is, for instance in the SWIR, larger than the variation observed among sun-exposed leaves sampled on one individual across the growing season (**dataset B; Fig. S13**), and would therefore negatively affect our ability to identify species trait variation. Besides, the calibration of such models seeking to harmonize spectra from different sources requires measurements of same samples using the involved spectroradiometers. Comparing measurements from databases consisting of independent datasets collected by several laboratories thus remains a challenge.

Jung et al. (2009) proposed an alternative approach using a white-reference dependent post-correction; the white references of different spectroradiometers are calibrated against a “master” white reference before comparing spectral measurements. Likewise, Pimstein et al. (2011) advocated the use of sand as an internal soil standard, being less expensive than a calibrated white reference (Spectralon®). Both approaches have been successfully adopted in the soil spectroscopy community, improving the retrieval of soil properties (e.g., clay content) by up to 10% (Romero, 2018). No equivalent procedure has so far been tested and adopted by the vegetation spectroscopy community. In our study, the measurement uncertainty associated with LOP was comparable to the uncertainty associated with a single measurement of standard fabrics with a low specular component (camouflage, cotton, plastic materials), suggesting that standard fabrics could be used as measurement standards to assess the measurement uncertainty associated with leaf reflectance. The uncertainty associated with repeated measurements of the latter materials fell below 0.006 and 0.02 reflectance units, representing between 4.5%-15% of leaf reflectance.

Similar to Hovi et al. (2018) and Potůčkováet al. (2016), we observed systematic differences in reflectance between leaf clip and integrating sphere sampling devices, while Lukěs et al. (2017) observed systematic differences among integrating spheres. Although Potůčková et al. (2016) successfully applied a linear model to homogenize spectral data measured with a leaf clip and an integrating sphere, our results suggest that we would not only need to develop a new model for each new combination of leaf sampling devices, but also for each new target. However, we observed the largest differences in leaf reflectance between leaf clip and integrating sphere measurements at specific wavelengths, generally used to retrieve pigment and water content. Prolonged exposure to the light beam due to successive measurements with a leaf clip or an integrating sphere may alter leaf chemical composition. By placing the leaf in slightly different positions for each measurement, we may also account for the natural variability in leaf chemical composition, and thus overestimate the actual differences in reflectance resulting from the use of different sampling devices. As Hovi et al. (2018) highlighted, correction procedures are challenging to develop, and care should be taken when comparing spectral data measured with different methods.

The integrating sphere appeared to produce more reproducible spectra with a low and stable measurement uncertainty, although the target was manually moved from one port to the other. The measurement uncertainty of integrating sphere measurements was independent of the optical properties of the target – at least regarding isotropy or anisotropy – and constant across a wide spectral range between 400-2500 nm. The integrating sphere has been the device of choice in numerous studies investigating LOP due to its reputation as a more stable measurement system (Jacquemoud and Baret, 1990; Milton, 2009). Nevertheless, we argue that spectral variation induced by the bi-directional measurement principle of a leaf clip provide important information on the diversity in LOP. Also, we advise to rotate the sample between measurements to better assess the effect of the specular component on the spectral variation.

Developing an uncertainty budget accounting for all sources of uncertainty and correcting systematic bias still appears a challenging task. Although in the proposed approach, we do not account for potential instability of the instrument over time, beyond applying a temperature correction (see **2.6**), or for specular properties, which increased the uncertainty associated with repeated measurements of fabrics, our results show that the calculation of the total uncertainty associated with a spectral measurement is a practical indicator of the measurement quality.

A more conservative approach would apply the measurement uncertainty associated with repeated measurements of standard fabrics to leaf spectral measurements. However, Anderson et al. (2011) highlighted that due to substantial differences between laboratory and field conditions, laboratory-derived uncertainties are not representative of uncertainties associated with field measurements. Thus we rather recommend to apply the developed approach to a given spectroradiometer-sampling device-target combination in field conditions. In addition, we suggest that measurements of a common standard (i.e.,

Spectralon or shared standard materials) should be adopted and systematically reported in metadata to enhance the comparability of spectral data in spectral libraries.

### 4.3. Partitioning of spectral variation into measurement uncertainty and biological variation

The largest spectral variation often corresponded to a maximum in leaf absorbance, which is driven by leaf morphology and chemistry. Also, spectral variation is a good indicator of morphological and physiological variation supporting the monitoring of species traits.

Spectral indices, building on the spectral variation between given wavelengths, and spectral diversity metrics using the variation across the full spectral range are the two main approaches commonly applied to predict leaf chemicals (Asner et al., 2009; Singh et al., 2015; Wang et al., 2020) and biodiversity (Schneider et al., 2017; Schweiger et al., 2018; Williams et al., 2020). However, a maximum in leaf absorbance corresponds to a minimum in reflectance; hence, the lower the reflectance, the lower the signal-to-noise ratio, and thus the larger the measurement uncertainty. Accounting for measurement uncertainty would therefore result in a more accurate assessment of biological variation. Yet, measurement uncertainty has not been systematically reported to date.

The spectral variation partitioning presented here allows for the distinction between measurement uncertainty and biological variation. To our knowledge, we show for the first time that measurement uncertainty represents roughly 2.5% of the spectral variation expected when sampling a tree over time (dataset B) or several trees in a forest site (datasets C and D). In addition, our measurements indicate that variation is at a minimum in spring and summer in the VNIR and SWIR spectral regions, respectively. Analyses conducted on a dataset including different levels of biological organization and phenology may thus provide more detailed insight into when in the growing season, and where in a spectrum, spectral measurements may best resolve species trait variation.

As already promoted by a handful of studies (Anderson at el., 2011; Hueni et al., 2017; Schaepman and Dangel, 2000), we thus advocate for a frequent calibration of field spectroradiometers, and a systematic characterization of uncertainty associated with spectral leaf measurements in field conditions. In addition, we suggest to partition spectral variation into measurement uncertainty and biological variation for a more accurate retrieval of LOP.

### 4.4. Spectral variation as a measure of species trait variation

Species trait variation describes the diversity in plant traits expressed by individuals within a species in response to their biotic and abiotic environment. We focused on species trait variation at three levels of biological organization: variation within an individual tree, and between individual trees sampled from single site and multiple sites. Within-tree variation occurs due to plasticity in relation to micro-environmental gradients within the crown (Atherton et al, 2017; Niimenets et al., 2014), phenology, and ontogeny (Albert, 2011), and at least in some cases to somatic mutations inherited from different meristems (Cruzan et al, 2020). We showed that out of an average biological variation of 40% in the SWIR, 30% is explained by the difference between shaded and sun-exposed leaves. This result appears to be in agreement with several studies investigating the effects of light gradients within the crown on leaf optical properties and reporting thicker leaves and lower water content in sun-exposed than in shaded leaves (Jacquemoud and Ustin, 2009, Niimenet et al. 2014). The remaining 10% of biological variation expressed in sun-exposed leaves was consistent across all datasets, suggesting that the biological variation captured by LOP represents the functional space allowed by the plasticity of sun-exposed leaves. We would like to note that our measurements also indicate larger within-tree variation in the fall. Similarly, Gao and Zhang (2006) observed a larger spectral diversity in fall and suggest that larger spectral variation allows for a better classification of salt marsh vegetation. Thus the time at which the largest within-individual variation is observed may also be suitable for observing the maximum between-individual variation, making a standardized approach even more important for comparing across individuals.

The biological variation between individuals from a unique or distinct population arises from the coexistence of genetically different individuals and plastic response to heterogenous environmental conditions (Albert, 2011). Spectral variation partitioning enables us to differentiate between within-individual versus between-individual variation. In our datasets, between-site variation appears larger than between-tree variation, which on average exceeds within-individual spectral variation when trees are measured within a short portion of the growing season (1 week). However, when considering the standard deviation within each biological level, variations within and among sites overlap in the visible range, while 95% confidence intervals of both levels were identical in the SWIR spectral range (**Fig. S12**). This finding echoes ecological studies showing a decreasing functional diversity between species when accounting for intraspecific variability of a single species (Ciancaruso, 2009; Violle, 2012) and suggests that intraspecific variability is an often-unmeasured indicator of how plants within a species fill a functional space (Schweiger, 2018). Thus the contribution of lower levels of biological organisation (branch, tree) to biological variation may exceed the contribution of higher levels of variation (e.g., between trees), perhaps especially in measurements of bulk properties such as spectroscopy. Neglecting the contribution of lower levels of organisation would overestimate the biological variation of the level of organisation of interest. This emphasizes the need for an appropriate characterization of lower levels of biological organisation which is often neglected in remote sensing studies.

Suitable sampling strategies can help capture species trait variability, and the biological variation retrieved from LOP allows for the monitoring of species traits at different levels of biological organization.

Additionally, it should be noted that spectral measurements are often carried out on harvested branches. Therefore, the measured variation can be an outcome of the leaf degradation proportional to the time between sample collection and spectral acquisition. In our studies, we did not account for this possible source of variation. Time between harvest and measurement was largest for dataset D, intermediate for dataset C, and least for dataset B (aside from the laboratory measurements associated with this dataset), due to different sampling approaches. We acknowledge that this can cause artifacts in derived measures of species trait variation. In our datasets, this may be an explanation for larger variation especially in the visible part of the spectrum in dataset D.

The analysis of the effect of sample size suggested that a minimum of 3 leaves sampled on 20 trees would give a good estimate of mean biological variation among sunlit canopy leaves of different trees in datasets that we investigated. This finding supports the sampling strategy adopted in the Swiss forest dataset, where each site corresponds to a collection of approximately 20 trees. In the same way, Petruzelli et al. (2017) suggested to use 4 leaves from 10 randomly-selected individuals to estimate the variation in specific leaf area (SLA) between individuals. However, an equally good characterization of the standard deviation (< ±5%) of between-individual variation would require a larger sample size of from 80 (in SWIR) to 110 (in VNIR) trees of an investigated population, again only if leaves are taken from a standardized position in the canopy across individuals (i.e., if substantial within-individual variation is ignored). Unlike Petruzelli’s study focusing on one specific trait, biological variation retrieved from LOP emerges from a range of plant traits. Traits commonly retrieved in the SWIR (e.g., water content, structural traits) appeared to require less sampling effort to characterize a mean value than do highly dynamic physiological traits (e.g., pigment content) expressed in the VNIR, but therefore a greater sampling effort to characterize the variation of individuals around the mean.

Field campaigns commonly adopt a selective sampling focusing on sun-exposed leaves preferentially located at the top of the canopy. This is justified by the fact that field campaigns often aim for comparability with spectral data from airborne sensors that can only measure the top of the canopy. Additionally, it ensures that the sampling is standardized across individuals. However, such selective sampling tends to underestimate the spectral diversity present within a population. In a biodiversity experiment, Schweiger et al. (2018) estimated that the spectral diversity at leaf level yielded a 10% better estimate of grassland productivity than tram-based remotely sensed spectra taken from above. Again, a more representative *in situ* sampling can help assess the fraction of variation that is either not accessible, or not resolved, by aerial spectra. To this end, a random sampling strategy irrespective of the light exposition in addition to the classic selective sampling – with a corresponding increase in the number of samples within individuals, as well as more individuals – would result in a better approximation of the commonly unmeasured variation (Petruzzeli et al, 2017; Violle et al., 2012).

## 5. Conclusion & Recommendations

Species trait variation is a key contributor to biodiversity and ecosystem function. The patterns of variation just within tree canopies likely supports different niches for habitation by the hundreds to thousands of other species that depend on any one tree species (Kennedy and Southwood, 1984), and variation among individuals is both an indicator of, and an explanation for, variation in environmental factors from nutrients and water, to species distributions (Salazar et al., 2017; Asner et al., 2017). We showed that leaf optical properties can indicate species trait variation at various biological and temporal scales. Partitioning of biological variation and measurement uncertainty helps disentangle the contribution of individual levels of biological organisation, from the individual to the forest, and to assess the potential biological information which can be retrieved at different spectral regions. We showed that relatively low measurement uncertainties for leaf reflectance given either low anisotropy, or an integrating sphere sampling device, permit the detection of variation at multiple levels of biological organization. However, the contribution of the measurement uncertainty to the spectral variation is not negligible, comprising 1-15% of total measured variation, and potentially overwhelming other sources of variation for leaves with high anisotropy. The characterization of the measurement uncertainty associated with leaf reflectance should consequently become a compulsory step during data processing, while sampling design should aim at minimizing it. We thus encourage the use of common measurement protocols and the systematic reporting of sufficient metadata in order to improve the traceability, the quality and comparability of spectral measurements. To this end, we have prepared a list of recommendations related to the instrument, the data acquisition, the sampling strategy, the data processing, and the metadata. The list aims to complement recommendations made in previous studies (Jiménez Michavila, 2015; Hueni et al., 2017;

Milton, 1987; Milton et al., 2009) with a focus on the use of sampling devices (i.e., leaf clip and integrating sphere).

- Regularly calibrate the spectroradiometer as wavelength drift may occur with time (Schaepman and Dangel, 2000)
- Regularly compare the white reference reflectance to the reflectance of a ‘pristine’ Spectralon reference panel as impurities and ultraviolet radiation may degrade the white reference over time.
- Establish the measurement uncertainty associated with the spectroradiometer and sampling device with a calibrated Spectralon reference panel and additional inert standard reference materials.
- Prefer repeated measurements at different spots of a target as prolongated exposure to the light beam may alter the chemical composition of biological material.
- Rotate the sample to minimize the impact of directional effects when measuring with a leaf clip device, but be aware that this will greatly increase uncertainty for highly anisotropic materials.
- Apply signal post-correction, including temperature correction to reduce measurement bias if necessary (Hueni, 2021).
- Estimate the uncertainty generated by the instrumentation, measurement protocols, and post-processing, and propagate these uncertainties to the resulting reflectance.
- Characterize the measurement uncertainty associated with the target, specifically if using a leaf clip (see 2.7) or any directional measurement device.
- Quantify the contribution of measurement uncertainty in relation to the total spectral variation (e.g., Table 1).
- Define the level of biological variation of interest and adopt a sampling strategy suitable to characterize and/or minimize the contribution of other levels of biological organisation to the biological variation, with specific attention to sampling bias and sample size.
- Random sampling produces a more faithful assessment of spectral variation. However, if vicarious calibration of airborne data is based on field measurements, sun-exposed leaves of the top of the canopy should be preferentially sampled. Random sampling within the canopy could be carried out in parallel to assess the fraction of variation that is not accessible to airborne sensors.
- Provide systematic metadata regarding all of the above considerations alongside your spectral data. The metadata should include the target description (e.g., species, sampling height, preferably coordinates), instrument characteristics (e.g., serial number), measurement protocol, calculation and correction procedures, and uncertainty budget.

Our study highlights the need for characterizing the contribution of various sources of uncertainty, such as the use of different spectroradiometers or different leaf clips of the same model, the repeatability of measurements in time, and the contamination of the white and black reference, to the measurement uncertainty associated with the leaf reflectance. In addition, increased sampling effort is required in order to better assess species trait variation from leaf reflectance. While experimental forests that are highly monitored and controlled may facilitate sampling, sampling in ’natural’ forests may become challenging, resource- and time-consuming. Innovative sampling strategies supported by machine learning may become an interesting alternative to better optimize target sampling efforts on the ground (Verrelst et al., 2020). Besides, statistical regressions (Singh et al., 2015), radiative transfer models (Féret et al., 2017, 2021), and large-scale comparisons (Meireles et al., 2020) allow for the retrieval of an increasing number of plant traits from leaf reflectance, providing additional insight into species traits deduced from spectral variation. Future work should focus on defining which meaningful species traits for biodiversity and ecosystem monitoring can be derived from leaf spectral measurements. While our approach can help better assess uncertainty associated with leaf spectral measurements, a broader and more accurate calibration of leaf spectral measurements is still to be done.

## Author contributions

CRediT taxonomy roles are listed with authors in alphabetical order. Conceptualization: F.P., G.G., M.C.S., M.E.S.; Project Administration: M.C.S.; Supervision: M.C.S., M.E.S., M.K.; Data Curation: E.A.C., F.P., G.G.; Formal Analysis: A.H., E.A.C., F.P., G.G.; Funding Acquisition and Resources: M.E.S., M.K.; Investigation: E.A.C., F.P., G.G.; Methodology: A.H., E.A.C., F.P., G.G., M.C.S., M.E.S., M.K.; Visualization: F.P.; Writing - Original Draft: E.A.C., F.P., G.G., M.C.S.; Writing - Review & Editing: A.H., E.A.C., F.P., G.G., M.C.S., M.E.S., M.K.

## Acknowledgments

The authors would like to thank Grün Stadt Zürich who allowed us to sample trees on their property, E. Magnanou and J. Garrigue who supported sampling in the French protected forest La Massane, and L. de Witte who supported the sampling in the Swiss forests as part of the long-term monitoring program of the Institute for Applied Plant Biology (IAP). We are grateful to H. Kühnle, A. Steppke and C. Li, who assisted with field spectroscopy measurements, and G. Wiesenberg and M.W.I. Schmidt, who supported the research project.

