## Supplementary material for "Variation in reflectance spectroscopy of European beech leaves captures phenology and biological hierarchies despite measurement uncertainties"

### Supplementary Data

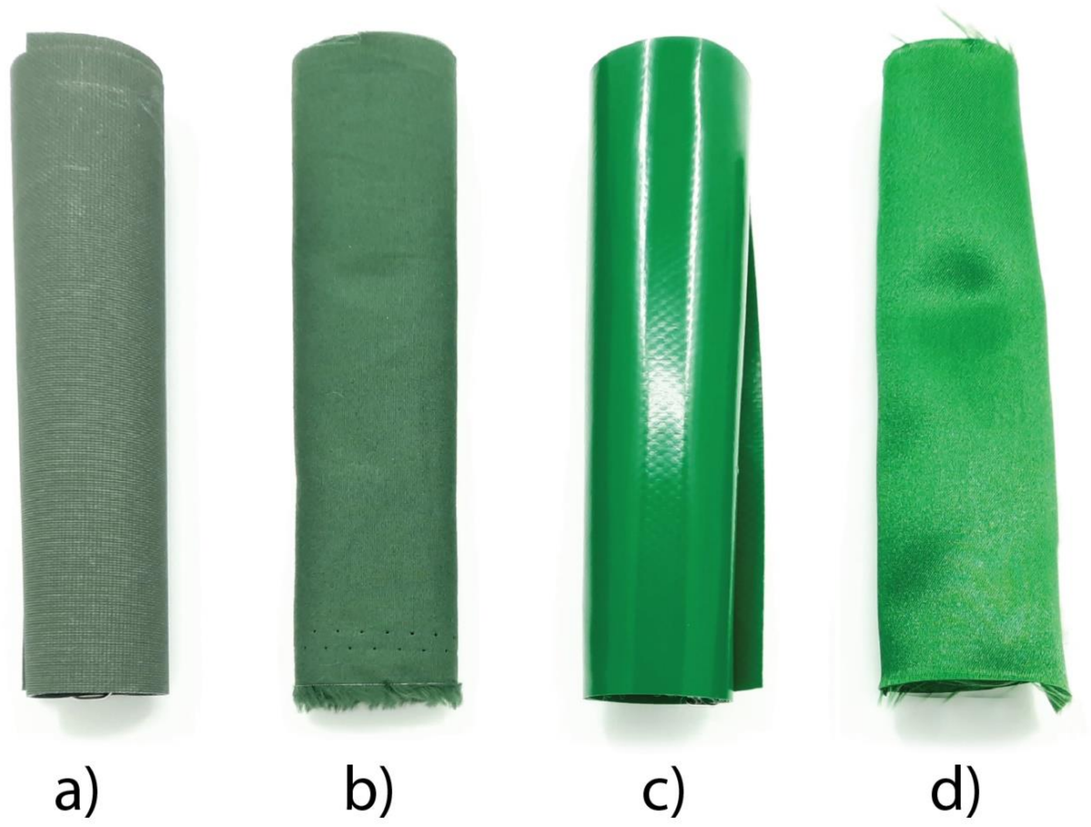


**Figure S1. Photographs and characteristics of the fabric standard materials: (a) camouflage fabric, (b) cotton fabric, (c) plastic fabric, (d) satin fabric.**

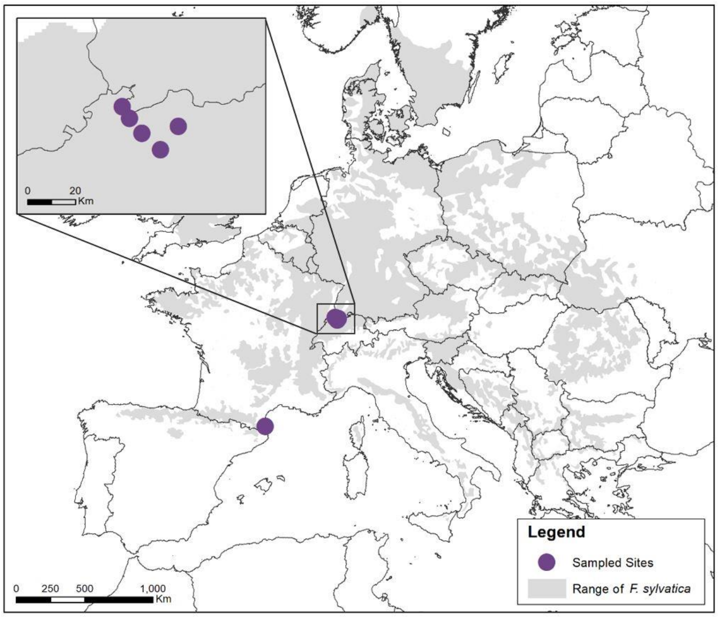


**Figure S2. Sampling sites of Dataset C and D, collected in La Massane reserve in the South of France and North-Eastern Switzerland, respectively. The sites are overlaid on the natural range of *F. sylvatica*.**

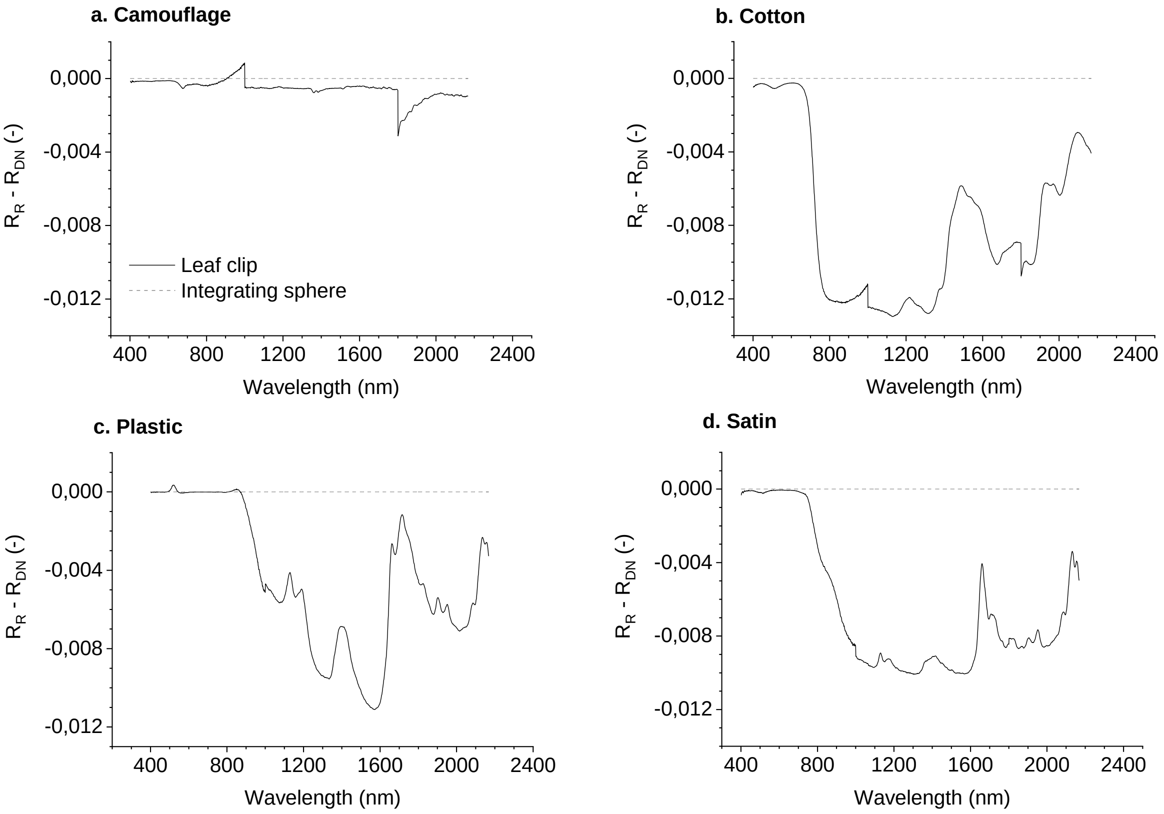


**Figure S3. Difference between the reflectance calculated from measurements performed with the leaf clip *(solid line)* and the integrating sphere *(dashed line)* in reflectance ($R_R$) and DN ($R_{DN}$) mode for four standard materials.** Note that the scale (0 to

-0.012 reflectance units) represents a maximum of approx. 2.2% of the calculated reflectance of these materials (see **Figure 2**).

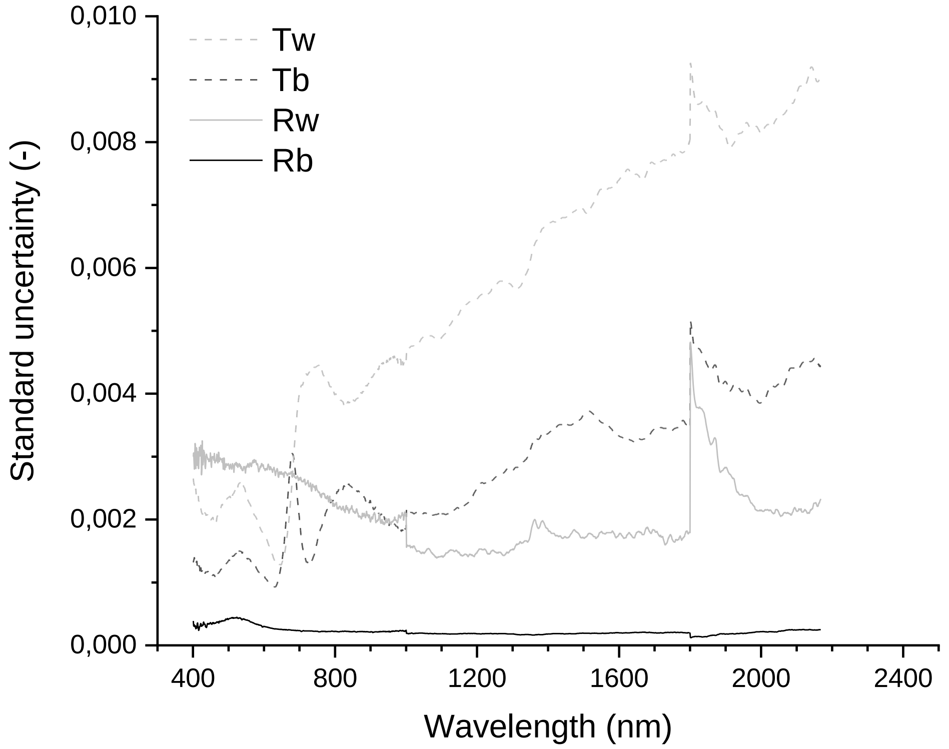


**Figure S4. Standard uncertainty of the four readings involved in the reflectance calculation**

The standard uncertainty corresponds to the standard deviation of each of the four measurement variables – white reference ($R_w$), white reference with target ($T_w$), black reference ($R_b$), black reference with target ($T_b$) - calculated from repeated measurements of fabrics (n=6).

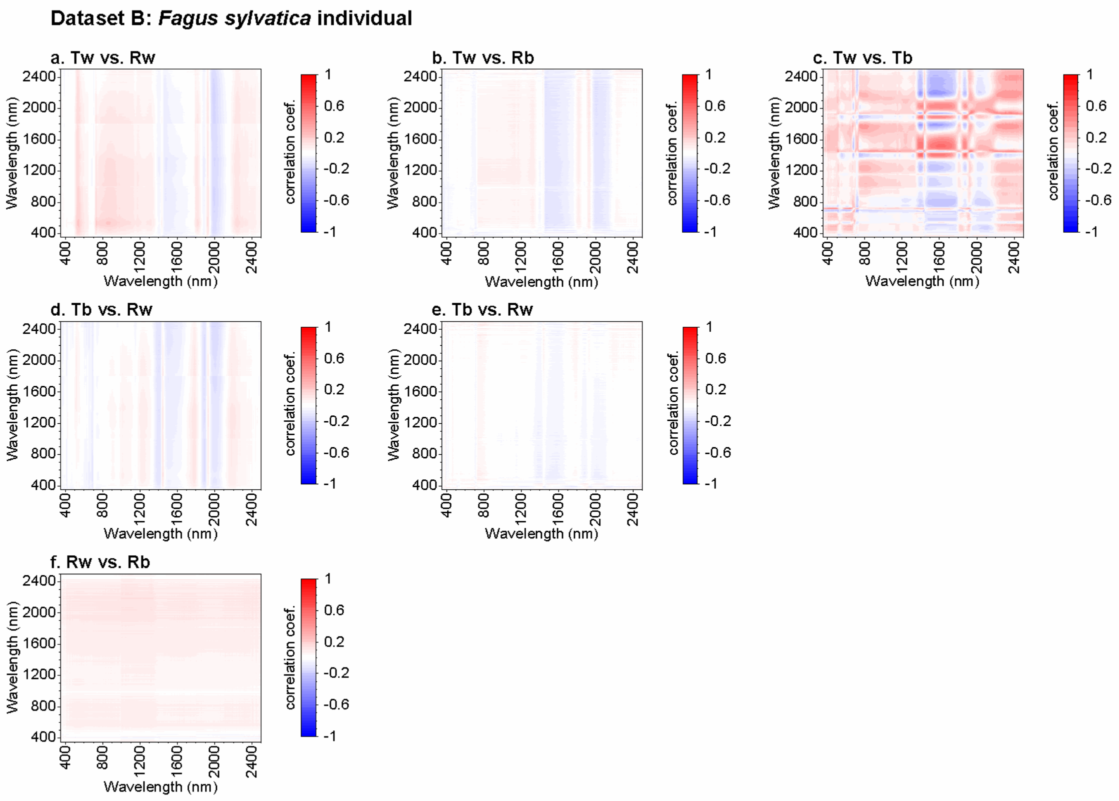


**Figure S5. Correlation between the standard uncertainty of readings in the dataset B**

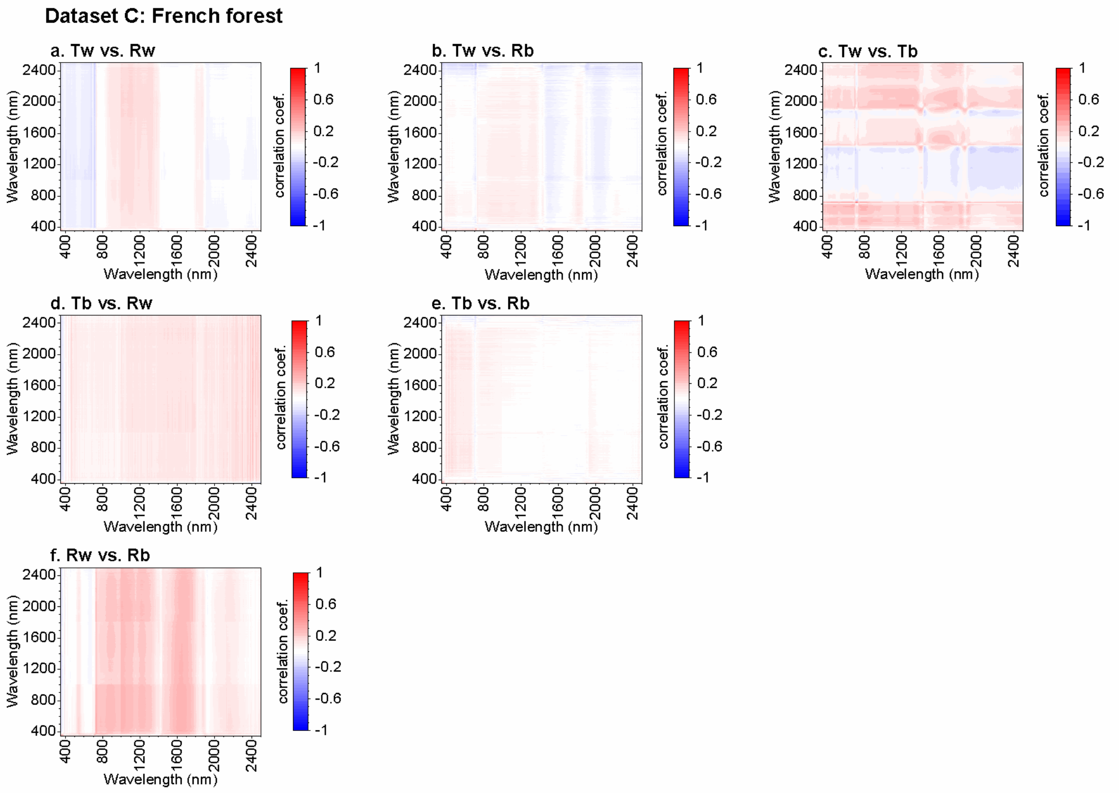

**Figure S6. Correlation between the standard uncertainty of readings in the dataset C**

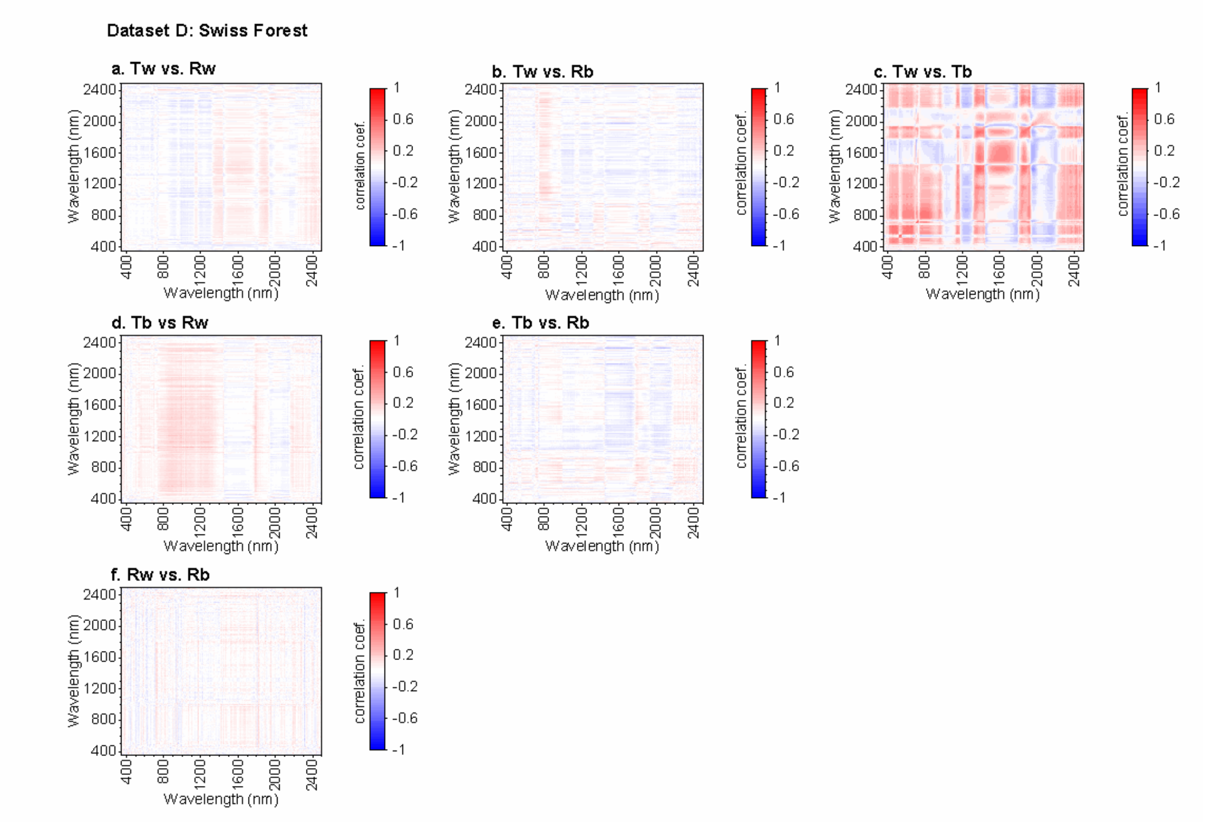


**Figure S7. Correlation between the standard uncertainty of readings in the dataset D**

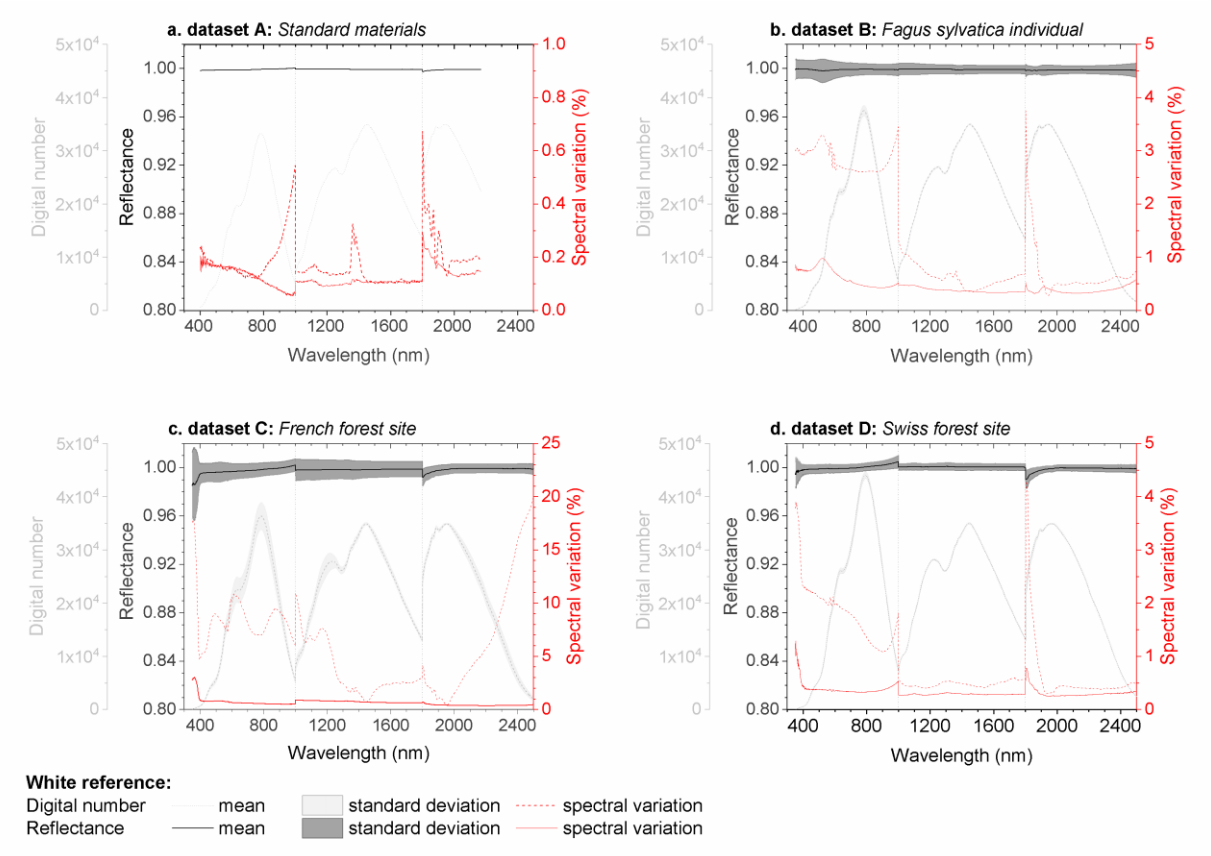


**Figure S8. Variation in the white reference standard of the leaf clip measured in digital numbers (DN) and reflectance mode, in laboratory (a) and field (b-d) conditions.**

Mean spectrum *(grey lines)*, standard deviation *(shaded area)* and coefficient of variation *(red lines)* of the white reference of the leaf clip measured in digital numbers *(dashed line),* and reflectance mode *(solid line)* without post-correction of the radiometric jumps.

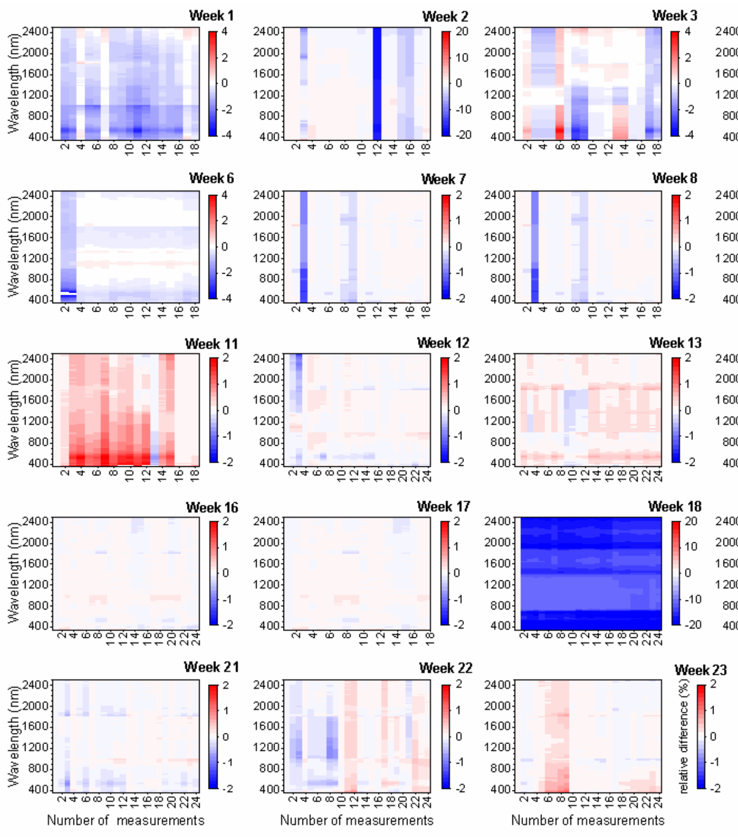

**Figure S9. Stability of the white reference over time.**

Each sampling week consisted in 18 to 24 measurements spread over 3 hours. The reflectance intensity of all the white reference measurements of a given week are compared to the first measurement of the white reference of the given week. Blue value indicates a lower reflectance and red a higher reflectance than the reflectance of the first white reference chosen as a reference. The scale is expressed in percentage.

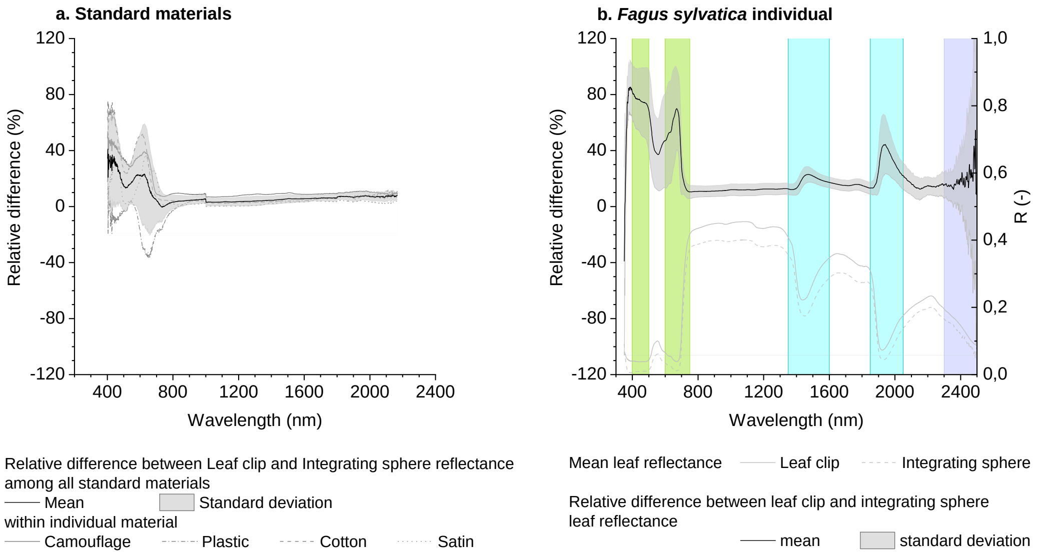


**Figure S10. Relative difference between the mean reflectance of four standard materials and *Fagus sylvatica* leaves successively measured with a leaf clip and an integrating sphere.** Crossed areas represent spectral ranges usually used to calculate spectral indices for pigments *(green),* water content *(blue)* and cellulose *(violet)*.

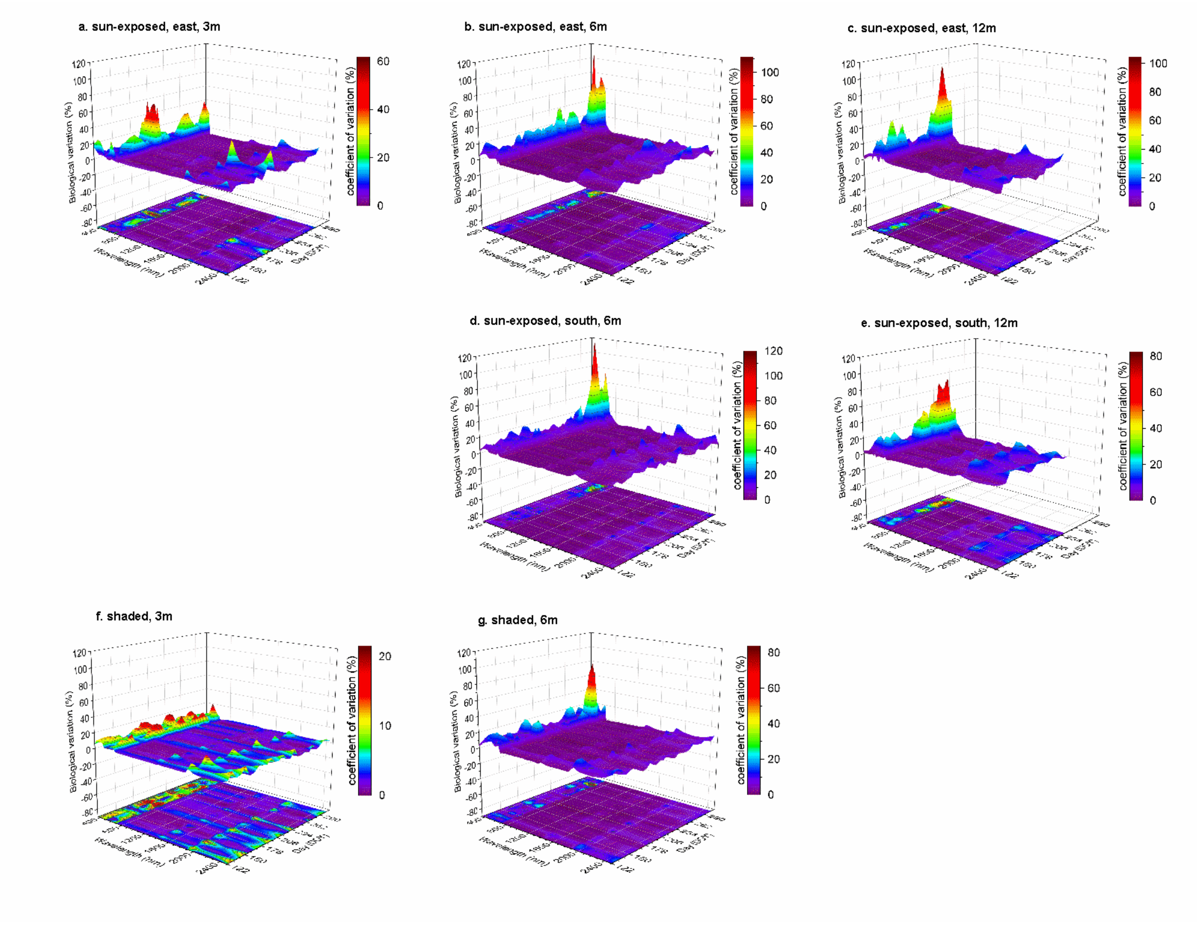


**Figure S11. Biological variation of several twigs of a *Fagus sylvatica* individual during the growing season.**

The biological variation describes the spectral diversity among leaves (n=3) belonging to the same twig. Sun-exposed leaves were weekly sampled from twigs located on the east-exposed side of the crown at (a) 3m, (b) 6m, and (c) 12m, and on the south-exposed side of the crown at (c) 6m and 12m. Shaded leaves were weekly sampled from twigs located under the crown at (f) 3m and (g) 6m.

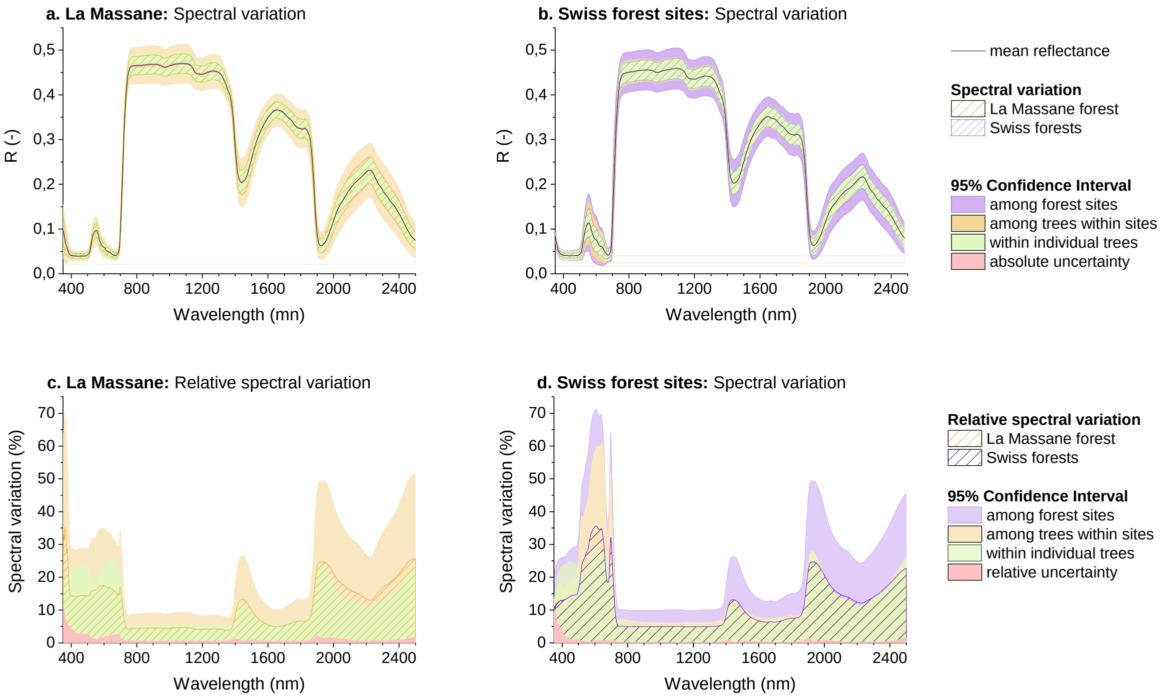


**Figure S12. 95% confidence interval of the spectral variation within a tree (green), between trees (orange) and between forest stands (purple) in comparison with the spectral variation observed in the dataset C (French forest) and D (Swiss forests) (crossed area).**

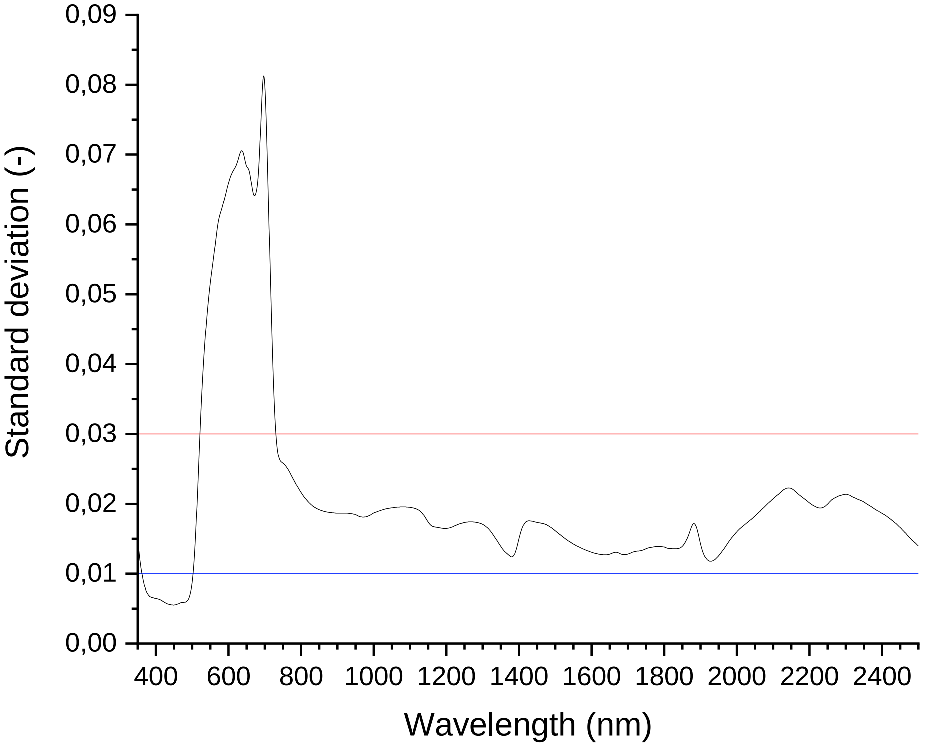


**Figure S13. Comparison of the spectral variation observed within an individual tree during the growing season (dataset B) with model errors of 0.1 (blue) and 0.3 (red) as used by Meireles et al. (2020) to harmonize spectra measured with different spectroradiometers.**

**Table S1**

| Sampling device | Data | Reflectance formula | Absolute uncertainty formula | Literature |
|---|---|---|---|---|
| Leaf clip | DN | $R = \frac{T_b}{R_w}$ | $U_{R,abs} = \sqrt{\left(\frac{1}{R_w}\right)^2 * \left(\frac{STD_{T_b}}{\sqrt{N}}\right)^2 + \left(\frac{-T_b}{{R_w}^2}\right)^2 * \left(\frac{STD_{R_w}}{\sqrt{N}}\right)^2}$ | (Potůčková, 2016; Hovi et al., 2018; Castro-Esau et al., 2006) |
| | reflectance | $R = \frac{T_b * R_w - T_w * R_b}{R_w - R_b}$ | $U_{R,abs} = \sqrt{\left(\frac{R_b * (T_w - T_b)}{(R_w - R_b)^2}\right)^2 * \left(\frac{STD_{R_w}}{\sqrt{N}}\right)^2 + \left(\frac{R_b}{R_w - R_b}\right)^2 * \left(\frac{STD_{T_w}}{\sqrt{N}}\right)^2 + \left(\frac{R_w * (T_w - T_b)}{(R_w - R_b)^2}\right)^2 * \left(\frac{STD_{R_b}}{\sqrt{N}}\right)^2 + \left(\frac{R_w}{R_w - R_b}\right)^2 * \left(\frac{STD_{T_b}}{\sqrt{N}}\right)^2}$ | (Miller et al., 1992) |
| Integrating sphere | DN | $R = \frac{I_s - I_d}{I_r - I_d} * R_r$ | $U_{R,abs} = \sqrt{\left(\frac{R_r}{I_r - I_d}\right)^2 * \left(\frac{STD_{I_s}}{\sqrt{N}}\right)^2 + \left(\frac{R_r * (I_s - I_r)}{(I_r - I_d)^2}\right)^2 * \left(\frac{STD_{I_d}}{\sqrt{N}}\right)^2 + \left(\frac{R_r * (I_d - I_s)}{(I_r - I_d)^2}\right)^2 * \left(\frac{STD_{I_r}}{\sqrt{N}}\right)^2}$ | (ASD manual) |
| | | $R = \left(\frac{I_s}{I_r} - S_r\right) * R_r$ | $U_{R,abs} = \sqrt{\left(\frac{R_r}{I_r}\right)^2 * \left(\frac{STD_{I_s}}{\sqrt{N}}\right)^2 + \left(\frac{-I_s * R_r}{{I_r}^2}\right)^2 * \left(\frac{STD_{I_r}}{\sqrt{N}}\right)^2 + (-R_r)^2 * \left(\frac{STD_{S_r}}{\sqrt{N}}\right)^2}$ | (Potůčková, 2016; Lukeš et al. 2018) |
| | reflectance | $R = \frac{I_s - I_d}{1 - I_d}$ | $U_{R,abs} = \sqrt{\left(\frac{R_r * (1 - I_d)}{(1 - I_s)^2}\right)^2 * \left(\frac{STD_{I_s}}{\sqrt{N}}\right)^2 + \left(\frac{R_r}{I_s - 1}\right)^2 * \left(\frac{STD_{I_d}}{\sqrt{N}}\right)^2}$ | (ASD manual) |

*Reflectance measurements with the leaf clip include four readings, the white reference ($R_w$), the white reference plus target ($T_w$), the black reference ($R_b$), and the black background standard plus target ($T_b$). Reflectance measurements with the integrating sphere include three readings, the reflectance reference ($I_r$), the reflectance sample ($I_s$) and the dark reading ($I_d$). $R_R$ corresponds to the nominal reflectance of the uncalibrated Spectralon® panel. $U_{R,abs}$ is the absolute uncertainty associated with the reflectance (R), while the standard uncertainty of each reading corresponds to the standard deviation among scans ($STD_x$) divided by the square root of the number of scans (N) recorded for a given reading.*
